# Single-cell transcriptomic analysis of skeletal muscle regeneration across mouse lifespan identifies altered stem cell states associated with senescence

**DOI:** 10.1101/2023.05.25.542370

**Authors:** Lauren D. Walter, Jessica L. Orton, Ern Hwei Hannah Fong, Viviana I. Maymi, Brian D. Rudd, Jennifer H. Elisseeff, Benjamin D. Cosgrove

## Abstract

Skeletal muscle regeneration is driven by the interaction of myogenic and non-myogenic cells. In aging, regeneration is impaired due to dysfunctions of myogenic and non-myogenic cells, but this is not understood comprehensively. We collected an integrated atlas of 273,923 single-cell transcriptomes from muscles of young, old, and geriatric mice (∼5, 20, 26 months-old) at six time-points following myotoxin injury. We identified eight cell types, including T and NK cells and macrophage subtypes, that displayed accelerated or delayed response dynamics between ages. Through pseudotime analysis, we observed myogenic cell states and trajectories specific to old and geriatric ages. To explain these age differences, we assessed cellular senescence by scoring experimentally derived and curated gene-lists. This pointed to an elevation of senescent-like subsets specifically within the self-renewing muscle stem cells in aged muscles. This resource provides a holistic portrait of the altered cellular states underlying skeletal muscle regenerative decline across mouse lifespan.

**EXTENDED SUMMARY:** Skeletal muscle regeneration relies on the orchestrated interaction of myogenic and non-myogenic cells with spatial and temporal coordination. The regenerative capacity of skeletal muscle declines with aging due to alterations in myogenic stem/progenitor cell states and functions, non-myogenic cell contributions, and systemic changes, all of which accrue with age. A holistic network-level view of the cell-intrinsic and -extrinsic changes influencing muscle stem/progenitor cell contributions to muscle regeneration across lifespan remains poorly resolved. To provide a comprehensive atlas of regenerative muscle cell states across mouse lifespan, we collected a compendium of 273,923 single-cell transcriptomes from hindlimb muscles of young, old, and geriatric (4-7, 20, and 26 months-old, respectively) mice at six closely sampled time-points following myotoxin injury. We identified 29 muscle-resident cell types, eight of which exhibited accelerated or delayed dynamics in their abundances between age groups, including T and NK cells and multiple macrophage subtypes, suggesting that the age-related decline in muscle repair may arise from temporal miscoordination of the inflammatory response. We performed a pseudotime analysis of myogenic cells across the regeneration timespan and found age-specific myogenic stem/progenitor cell trajectories in old and geriatric muscles. Given the critical role that cellular senescence plays in limiting cell contributions in aged tissues, we built a series of tools to bioinformatically identify senescence in these single-cell data and assess their ability to identify senescence within key myogenic stages. By comparing single-cell senescence scores to co-expression of hallmark senescence genes *Cdkn2a* and *Cdkn1a*, we found that an experimentally derived gene-list derived from a muscle foreign body response (FBR) fibrosis model accurately (receiver-operator curve AUC = 0.82-0.86) identified senescent-like myogenic cells across mouse ages, injury time-points, and cell-cycle states, in a manner comparable to curated gene-lists. Further, this scoring approach pinpointed transitory senescence subsets within the myogenic stem/progenitor cell trajectory that are related to stalled MuSC self-renewal states across all ages of mice. This new resource of mouse skeletal muscle aging provides a comprehensive portrait of the changing cellular states and interaction network underlying skeletal muscle regeneration across mouse lifespan.

## INTRODUCTION

Skeletal muscle is heterogeneously composed of interacting immune, stromal, and myogenic cells that contribute to the maintenance and regeneration of muscle by regulating muscle stem cell (MuSC) quiescence, proliferation, and differentiation.^1^ MuSCs are found between the basal lamina and the plasma membrane of myofibers and are essential for the initial development of muscle and in muscle regeneration.^2, 3^ The MuSC population is maintained across multiple cycles of growth and regeneration by asymmetrical division which generates additional MuSCs and *Myod1*+ myoblasts.^2, 3^ Myoblasts further expand, differentiate, and fuse to form new *Myog*+ myocytes.^2, 3^ The paired box protein 7 (*Pax7*) transcription factor, predominantly expressed in MuSCs, regulates the expression of myogenic regulatory factors such as *Myf5* and *Myod1*.^3^

During homeostasis, the skeletal muscle microenvironment maintains signals to keep MuSCs in a non-cycling state and resident mast cells and macrophages monitor for damage.^1, 4^ Following muscle injuries, the mature myofibers undergo necrosis and the individual myonuclei undergo apoptosis.^5, 6^ Once damage is detected by resident immune cells, inflammatory cells like neutrophils and macrophages are recruited to the damaged site.^1, 2^ First, resident macrophages release *Cxcl1* and *Ccl2*, neutrophil chemoattractants, to signal for neutrophils to invade.^1, 4^ Neutrophils reach their maximum abundance in the damaged muscle between 12-24 hours post-injury, after which they quickly return to basal levels.^4^ Resident Cd8+ T cells also respond early to muscle injury by producing *Ccl2* and recruiting macrophages to the injured muscle.^4^ Circulating monocytes and macrophages enter the muscle environment which is enriched with pro-inflammatory cytokines that activate macrophages.^1, 4^ These activated macrophages clear cellular debris and promote myogenic cell proliferation.^1, 5^ The active phagocytic macrophages peak in abundance at 2 days post-injury (dpi) and they are replaced by non-phagocytic macrophages that peak in abundance at 4-7 dpi.^4^ The non-phagocytic macrophages help maintain myogenic cell differentiation, resolve inflammation, and support the production of connective tissue.^1, 4, 5^ If macrophages do not clear cellular debris and promote myogenic cell proliferation and differentiation, the muscle remains inflamed and there are repeated cycles of necrosis and regeneration.^5^ The damaged myofibers are then replaced with adipose tissue, fibrotic tissue, or bone, instead of new myofibers.^5^ A prior study found that there was an increase in the number of anti-inflammatory macrophages which can contribute to an increase in fibrosis in aged muscle.^7^ It remains controversial how monocytes and macrophages should be classified and whether broad pro- and anti-inflammatory definitions should be used or whether more varied phenotypic identities better capture their molecular and functional plasticity.^8^ In this work, we sought to distinguish monocyte/macrophage populations by markers and by resident status.

Prior studies have found that both CD8+ and CD4+, especially FoxP3+ regulatory T cells, infiltrate and help repair damaged muscle.^9^ In aging, there is a decline in new naive T cells released from the Thymus.^10^ To compensate for this loss, CD4+ T cells in the periphery proliferate resulting in a shift towards more memory T cell populations throughout aging.^9^ Within the memory T cell population, an increase in senescent and exhausted T cells is observed in aging.^9^

In addition to resident immune cells, fibro-adipogenic progenitors (FAPs), bipotent progenitor cells that can differentiate into fibroblasts and adipocytes, are a part of the skeletal muscle microenvironment.^1^ FAPs are quiescent during homeostasis and activate upon injury.^4^ There is some evidence that in the early stages of injury response FAPs control immune infiltration and in later stages FAPs control muscle remodeling.^11^ They reach peak abundance at 3 dpi and return to homeostatic levels by 14 dpi.^4^ The expansion and decline of FAPs is regulated by myeloid cells and FAPs that are activated to the fibrogenic phenotype are regulated by anti-inflammatory macrophages.^4^ Fibrogenic FAPs are the primary producers of connective tissue in injured muscle.^4^

Multiple factors that contribute to the reduced functionality of MuSCs in aged tissues have been reported, including an excess of FAPs and fibroblasts, misbalanced cell division, and the establishment of senescent MuSCs.^2, 4, 12, 13^ Senescence is characterized by a combination of hallmarks. These include prolonged DNA damage response activation, upregulation of the cell cycle inhibitors p16^INK4a^ (encoded by *Cdkn2a* and referred to as p16 hereafter) and p21^Cip^^1^ (encoded by *Cdkn1a* and referred to as p21 hereafter) and anti-apoptotic BCL-2 proteins, an increase in reactive oxygen species levels and of senescence-associated-β-galactosidase (SA-β-gal), and a senescence-associated secretory phenotype (SASP).^14^ Prior studies have used the cell cycle proteins p16, p21, p53, and Rb to differentiate between dividing and non-diving cells, but the non-diving cells can include senescent cells and quiescent cells.^15^ SA-β-gal is also commonly used to identify senescent cells, but it is also detected in quiescent cells and in stressed cells.^15^ Because these markers are not unique to senescent cells and because senescent cells are heterogeneous, it has been challenging to identify biomarkers that can accurately and consistently identify senescent cells.^16^

There are extrinsic changes, such as an increase in FAPs, and intrinsic factors, such as a reduction in asymmetrical self-renewal and an increase in senescent MuSCs, that disrupt skeletal muscle homeostasis and regeneration in aging.^2^ Aged MuSCs exhibit a decline in self-renewal and ability to differentiate, thus reducing the MuSC pool.^2, 4, 12, 13^ Compared to young MuSCs, fewer aged MuSCs are found in quiescence, due to either elevated activation or entering a pre-senescent state.^12, 13^ It remains unclear how these extrinsic and intrinsic changes are integrated systematically and how heterogeneities related to cellular senescence within both myogenic and non-myogenic cell populations contribute, in part due to a paucity of holistic analyses of these alterations. Further, it has been posited that temporal and spatial discoordination between the dynamics in key cell types during the repair process leads to inefficient outcomes.^2^

Single-cell methods have been used previously to understand skeletal muscle homeostasis and regeneration at various ages.^11, 17–32^ Recent reports have provided insights into MuSC dysfunction with aging. A recent human skeletal muscle study observed a decline in the proportion of MuSCs with age and that *IGFN1*, which is needed for myoblast fusion and differentiation, is decreased in old (∼75 years old) muscle.^28^ This report found no difference in *Cdkn2a* expression by age, but did identify a senescent myonuclei population that expressed *Cdkn1a* that was more frequent in old than young human samples.^28^ A recent study on mouse skeletal muscle aging based on mass cytometry observed that CD47^hi^ MuSCs occur with a higher frequency in aged mice and exhibit poor regenerative capacity.^33^ Another report demonstrated that quiescent MuSCs in aged mice have reduced expression of Cyclin D1, which is needed for proper MuSC activation and muscle regeneration.^34^

To evaluate the factors that contribute to the age-related decline in skeletal muscle regeneration in a more comprehensive manner, we have generated a new single-cell RNA sequencing (scRNA-seq) analysis of uninjured (day 0) and myotoxin-injured (days 1, 2, 3.5, 5, and 7) tibialis anterior (TA) muscles from young, old, and geriatric mice. We identified a total of 29 cell types, 8 of which had a significant difference in their abundances throughout regeneration by age. We confirmed changes in age-specific T cell abundance by immunohistochemistry and flow cytometry. Given the role that cellular senescence plays in limiting cell contributions in aged tissues, we tested a series of tools to bioinformatically identify senescence in these single-cell data and found a transfer-learning based scoring approach accurately classified senescent-like myogenic cells across ages and cell cycling states. This scoring approach revealed that senescent-like subsets exist at key transitional self-renewal states within the myogenic stem/progenitor cell pseudotime trajectory across all ages but are elevated in aged muscles. This resource of mouse skeletal muscle aging provides a more comprehensive portrait of the changing cellular states underlying skeletal muscle regeneration across mouse lifespan.

## RESULTS

### Single-cell RNA-sequencing analysis of skeletal muscle regeneration across mouse lifespan

To comprehensively evaluate skeletal muscle homeostasis and regeneration throughout aging, we performed single-cell RNA sequencing (scRNA-seq) on 65 mouse skeletal muscle samples with the 10x Chromium v2 and v3 platforms. Muscle damage was induced in young (4-7 months-old [mo]), old (20 mo), and geriatric (26 mo) C57BL/6J mice by injecting the tibialis anterior (TA) muscles with notexin. The injured and uninjured muscles were collected at six time points (days 0, 1, 2, 3.5, 5, and 7) (**Figure 1A, B**). Together, these 65 scRNA-seq samples, newly reported here and from our two prior reports^20, 35^, contained a total of 365,710 cell barcodes prior to quality control and filtering (**Supplementary Figure 1A**, **Extended Data File 1**). All samples were processed by aligning sequencing reads to the mm10 mouse reference genome, removing ambient RNA signatures with SoupX^36^, removing low quality cells, and identifying and removing doublets with DoubletFinder^37^ (**Figure 1C**, **Supplementary Figure 1**). The samples were integrated with Harmony^38^ to correct for batch effects (**Figure 1C**). The final dataset contained 273,923 cells. We observed that the cell number was relatively evenly distributed across the three age groups (**Figure 1D,G**), and within each age group, the number of cells from each time point was relatively consistent (**Figure 1E**).

**Figure 1:**
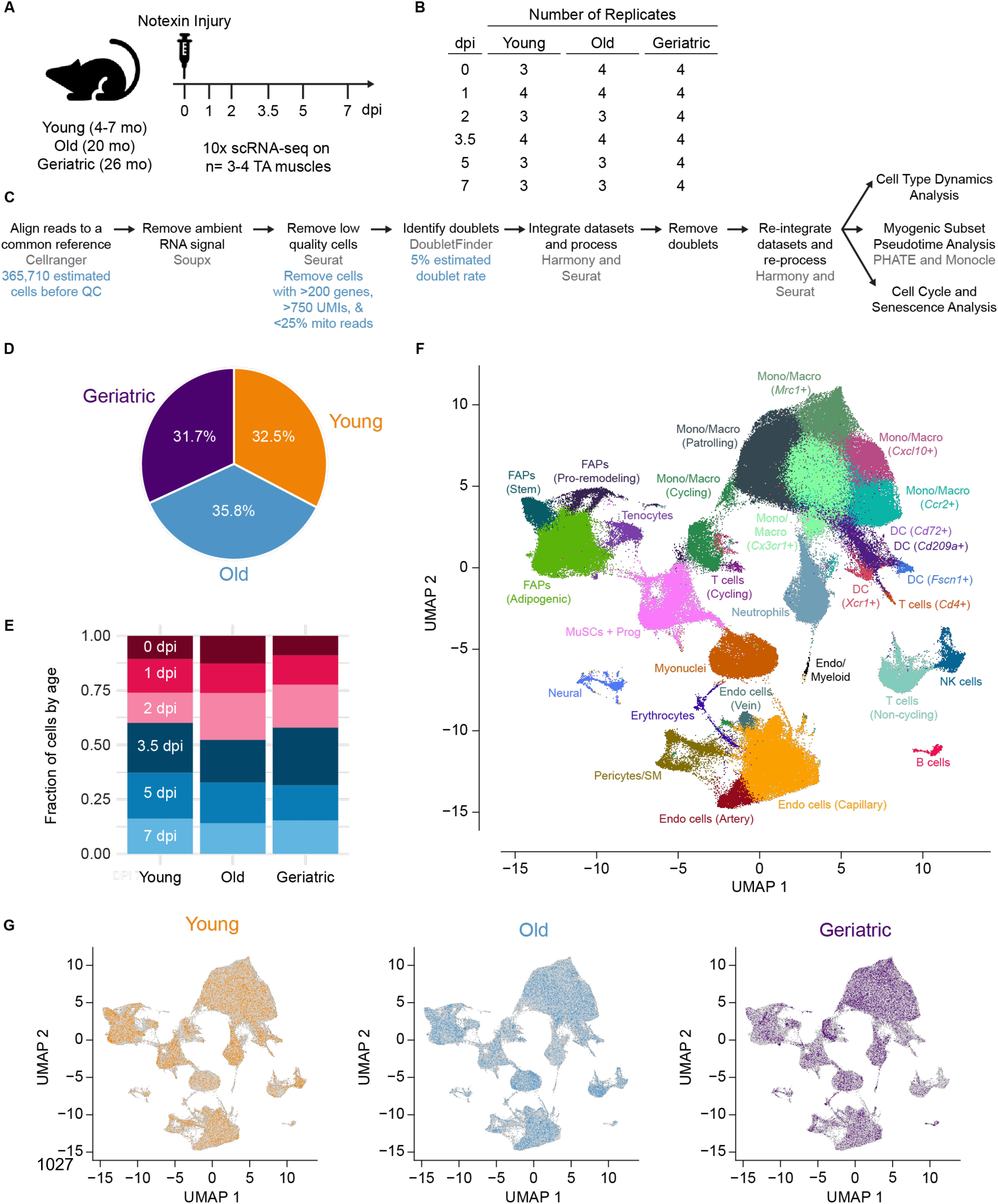
Assembly of scRNA-seq atlas of skeletal muscle regeneration across mouse aging. (**A-B**) Overview of experimental design. 3’ scRNA-seq (10X Chromium v2 and v3) was performed on dissociated tibialis anterior (TA) muscles from young (4-7 mo), old (20 mo), and geriatric (26 mo) mice (both sexes) 0-7 days post-notexin injury (dpi) with n = 3-4 replicates per age and dpi (**B**). (**C**) Processing workflow. Each scRNA-seq sample was aligned to the mm10 mouse reference genome, ambient RNA was removed by SoupX, low quality cells were identified and removed, and doublets were identified and removed. All samples were then integrated with Harmony, resulting in a final dataset containing 273,923 cells from 65 samples. See **Supplementary** Figure 1 and **Extended Data File 1** for additional detail. (**D**) Fraction of cells from each age group. (**E**) Fraction of cells from each dpi within each age group. (**F**-**G**) UMAP representations of the final dataset. Cells colored by manually assigned cell type IDs based on the expression of hallmark skeletal muscle genes (see **Supplementary** Figures 2 and 3) (**F**). Cells are colored by age group, with all other cells in gray (**G**).

### Multi-step identification of diverse cell types by clustering

After data integration, shared nearest neighbor (SNN) clustering was performed, and canonical marker genes were used to manually identify cell types. The initial clustering resulted in 24 clusters that each received a unique cell type annotation. Of these, nine were myeloid cell clusters (monocytes, macrophages, dendritic cells, and neutrophils) that exhibited similar expression profiles. To further clarify the myeloid cells found in the final dataset, the 9 myeloid cell clusters were subset out, re-clustered, and re-embedded (**Supplementary Figure 2A-C**). This resulted in 15 clusters that each received a unique cell type annotation based on known myeloid markers (**Supplementary Figure 2C-D**). Although this re-clustering did not further clarify the monocyte and macrophage annotations, it did help to identify more specific dendritic cell and T cell subtypes (**Supplementary Figure 2C-D**). The dendritic cell and T cell subtype annotations were transferred from the myeloid subset back to the final dataset based on the cell barcode. The monocyte and macrophage annotations were not changed based on the myeloid subset. With the additional myeloid subset annotations, we identified 29 distinct cell types in the final dataset (**Figure 1F**, **Supplementary Figure 3**). Compared to prior scRNA-seq and snRNA-seq skeletal muscle studies we identified similar broad cell types and we identified more specific endothelial, FAP, and immune cell subsets.^11, 18–30^

Lymphoid cell types were 5 of the 17 immune cell clusters. We identified a rare B cell cluster (0.47% of the final dataset) that expressed *Cd19*, *Cd22*, and *Ms4a1* (**Supplementary Figure 3A**). We identified a natural killer (NK) cell cluster that expressed *Nkg7*, *Gzma*, *Klra4*, and *Klre1* but, importantly, did not express T cell markers (**Supplementary Figure 3A**). We identified three T cell clusters all of which expressed *Cd8a* and *Cd8b1*. One of the T cell populations more strongly expressed *Cd4* while the other two T cell populations expressed *Cd3e*. One of the *Cd3e*+ clusters also strongly expressed the cycling markers *Cdk1* and *Hmgb2* and were thus identified as cycling *Cd3e*+ T cells (**Supplementary Figure 3A**). The other *Cd3e*+ T cell population appears to be non-cycling. The B cells, NK cells, and the non-cycling *Cd3e*+ T cells were identified as unique clusters in the initial clustering of the final dataset. The *Cd4*+ and cycling *Cd3e*+ T cells were identified when we subset and re-clustered the myeloid populations (**Supplementary Figures 2C-D** and **3A**).

Although we incubated the single-cell suspension in erythrocyte lysis buffer, we did see a small (0.42% of the final dataset) erythrocyte cluster that uniquely and strongly expressed a variety of hemoglobin genes, including *Hba-a1* and *Hbb-bs* (**Supplementary Figure 3A**). Erythrocytes are not a native cell type in skeletal muscle, so we have excluded them from the cell type dynamics analysis.

We identified three FAPs populations (adipogenic, pro-remodeling, and stem), a tenocytes population, and a Schwann and neural/glial cell population. All three of the FAPs populations expressed *Pdgfra* and *Col3a1* (**Supplementary Figure 3B**). The adipogenic FAPs also expressed *Adam12*, *Bmb5*, *Myoc*, *Col1a1*, *Dcn*, *Mmp2*, and *Apod*. The pro-remodeling FAPs uniquely expressed cycling genes like *Cdk1* and *Tyms* in addition to other FAPs markers like *Tnfalp6*, *Il33*, *Adam12*, *Bgn*, and *Hdlbp*. The stem FAPs also expressed *Igfbp5*, *Dpp4*, *Cd34*, *Gsn*, and *Mmp2*. The tenocyte population expressed some FAPs markers like *Col1a1*, *Dcn*, and *Apod* and they expressed tenocyte-specific markers *Tnmd* and *Scx*. The Schwann and neural/glial cell cluster expressed *Ptn* and *Mpz* (**Supplementary Figure 3B**).

We identified a pericytes and smooth muscle cells cluster which expressed the pericyte-specific gene *Rgs5* and *Acta2*, *Myl9*, and *Myh11* (**Supplementary Figure 3C**).^39^ Four endothelial clusters were identified, and they shared strong expression of *Cdh5* and *Pecam1* (**Supplementary Figure 3C**). The arterial endothelial cells uniquely expressed *Alpl* and *Hey1*, the capillary endothelial cells strongly expressed *Lpl*, and the venous endothelial cells expressed *Vwf*, *Hlf1a*, *Icam1*, *Lrg1*, and *Aplnr* (**Supplementary Figure 3C**). The fourth endothelial cluster expressed endothelial markers like *Cdh5* and *Pecam1* and myeloid markers like *S100a8/9*, *Csf1*, and *Itgam* (**Supplementary Figure 3C**). This cluster was small (0.073% of the final dataset) and was made up of cells from multiple replicates from the three ages at 0-2 dpi (**Supplementary Figure 5E**). Because this cluster was not unique to a single replicate, or single age, or single time point, we have maintained it in our analysis.

### Comparison of cell type dynamics in skeletal muscle regeneration across mouse ages

For each time point independent of age, we calculated the percent of cells from each cell type. In agreement with previous findings, uninjured skeletal muscle (day 0) was mainly composed of endothelial cells (49.2%), FAPs (20.4%), and myonuclei (15.1%) and there were small populations of immune cells (2.7%) and MuSCs and progenitors (2.0%) (**Supplementary Figure 4A**, **Supplementary Table 1A**).^20, 40^ Of the endothelial cells at day 0, the most prominent subtype were the capillary endothelial cells (42.4%) (**Supplementary Figure 4B**, **Supplementary Table 1B**). The FAPs present at day 0 were mainly adipogenic FAPs (15.6%) (**Supplementary Figure 4B**, **Supplementary Table 1B**). Of the immune cells present at day 0, there were a variety of monocytes and macrophages (47.2%), B cells (14.6%), non-cycling *Cd3e*+ T cells (11.6%), *Cd209a*+ Dendritic cells (10.6%), and Neutrophils (9.3%) (**Supplementary Figure 4D**, **Supplementary Table 1D**).^40^

Following injury (days 1, 2, 3.5, and 5), the most abundant general cell type was the immune cells (60.9%, 79.7%, 75.2%, 63.2%, respectively) (**Figure 2K**, **Supplementary Figure 4B**, **Supplementary Table 1B**). As in previous studies^20, 40^, non-immune cells like endothelial cells, FAPs, and myonuclei were present following injury, but at transiently lower relative abundances. As expected, the most abundant immune cells immediately after injury (day 1) were neutrophils (32.0%), *Ccr2*+ monocytes/macrophages (19.7%), and *Ctsa*+ patrolling monocytes/macrophages (13.7%) (**Supplementary Figure 4C**, **Supplementary Table 1C**).^20, 40^ Immediately following injury (days 1 and 2), there was a more pro-inflammatory environment, as evident by the abundance of *Ccr2*+ monocytes/macrophages. This was followed by a shift at days 3.5 and 5 to a more anti-inflammatory cell population, as evident by the peak in abundance of *Cx3cr1*+ monocytes/macrophages at day 5 (28.7%) (**Supplementary Figure 4C**, **Supplementary Table 1C**).^20, 40^ By day 7 the cell type abundances were returning to the abundances observed at day 0, but there was still a substantial immune cell population (32.1%) (**Supplementary Figure 4A**, **Supplementary Table 1A**). The immune population at day 7 mainly consists of *Cx3cr1*+ monocytes/macrophages (24.1%), *Ctsa*+ patrolling monocytes/macrophages (13.2%), and non-cycling *Cd3e*+ T cells (17.8%) (**Supplementary Figure 4D**, **Supplementary Table 1D**).

**Figure 2:**
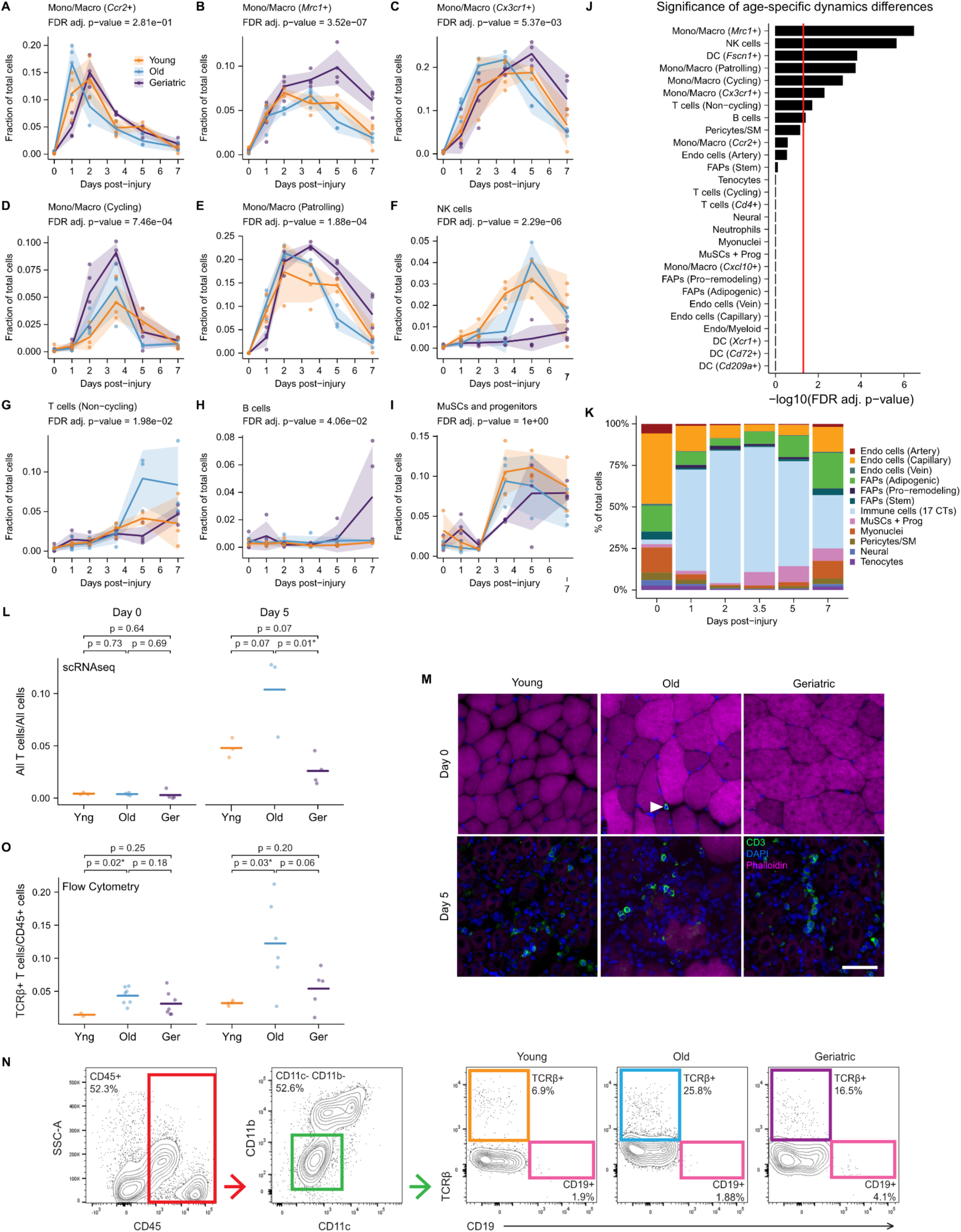
Age-related changes to cell dynamics during skeletal muscle regeneration. (**A**-**I**) Line plots showing cell type relative abundance as a fraction of total cells from 0-7 days post injury (dpi). For each sample, the number of cells of the reported type was divided by the total number of cells (excluding erythrocytes). Points are each sample (n = 3-4). Ribbon is the standard deviation. Statistical significance of age-specific cell type dynamics was evaluated using non-linear modeling and FDR-corrected p-values are reported (see **Supplementary** Figure 6). (**J**) Statistical significance of age-specific cell type dynamics differences as reported by FDR-corrected p-values from non-linear modeling (see **Supplementary** Figure 6). Red line denotes the FDR-adjusted p = 0.05 threshold. (**K**) Stacked bar plot of the fraction of cell types across all dpis. All 17 immune cell clusters were grouped into a unified “Immune cells (17 CTs)” cluster, which is separated out in **Supplementary** Figure 4. (**L**) Scatter plots of the fraction of all T cells (3 CTs) at 0 and 5 dpi from the scRNA-seq data. For each sample, the number of all T cells (3 CTs) was divided by the total number of cells (excluding erythrocytes). Points are each sample (n = 3-4). Line is the mean for each age group. Significance was evaluated using the Student’s t-test. (**M**) Immunohistochemical analysis of CD3+ T cells (green) at day 0 and day 5 post-injury in tibialis anterior (TA) muscles from young (5 mo), old (20 mo), and geriatric (26 mo) mice with DAPI (blue) as a nuclear counterstain and Phalloidin-750 (pink) as myofiber counterstain. Scale bar, 50 µm. Arrow denotes CD3+ T cell. (**N**) Flow cytometric analysis of TCRβ+ T cells. After gating single viable cells by FSC/SSC and a fixable viability dye (not shown), CD45+ hematopoietic cells, CD11b-CD11c-non-myeloid cells, and TCRβ+ CD19-T cells were gated sequentially. (**O**) Scatter plots of the fraction of TCRβ+ cells out of all CD45+ cells at 0 and 5 dpi. Points are each sample (n = 2-7). Line is the mean for each age group. Significance was evaluated using the Student’s t-test.

We next conducted an analysis of cell-type dynamics across age groups by comparing the abundance of a given cell type over the entire injury time course. Of the 28 cell types, eight were identified as having significantly different cell type dynamics across the three ages using a non-linear modeling approach with multiple hypotheses correction (**Figure 2J**, **Supplementary Table 2**). In response to injury, we first observed an increase in neutrophils which peaked in abundance at day 1 and returned to day 0 levels by day 3.5 (**Supplementary Figure 5N**). We also observed a peak in the abundance of the *Ccr2*+ monocytes/macrophages at days 1 and 2 (**Figure 2A**) while the cycling monocytes/macrophages peaked in abundance at day 3.5 (**Figure 2D**) and the *Cx3cr1*+ monocytes/macrophages peaked in abundance at days 3.5 and 5 (**Figure 2C**). We observed two monocyte/macrophage populations, *Mrc1*+ and *Ctsa*+ monocytes/macrophages, that responded early to injury (day 2) and remained high through day 5 (**Figure 2B,E**). The geriatric *Mrc1*+ monocytes/macrophages maintained a higher abundance from day 2 to day 7 compared to the young and old cells (**Figure 2B,J**). Additionally, some lymphoid cell types like NK cells, *Cd3e*+ non-cycling T cells, and B cells started to increase in abundance at day 2, day 3.5, and day 5, respectively (**Figure 2F-H**). We observed a similar pattern when looking at all three T cell populations combined (**Supplementary Figure 5T**). The geriatric NK cells did not increase in abundance within the 7-day time course, unlike the young and old cells (**Figure 2F**).

When looking at all three T cell populations combined, we detected very few T cells and no age-specific differences in abundance at day 0. However, at day 5 we observed a higher abundance of T cells in old samples, with a significant difference between the old and geriatric samples (**Figure 2L**; Student’s t-test, p-value = 0.01*). To confirm this, we performed immunohistochemistry on sectioned TA muscles and observed that CD3+ T cells are detected more abundantly at day 5 compared to day 0 in the TAs of young, old, and geriatric mice (**Figure 2M**). We further used flow cytometry to quantify CD45+CD11c–CD11b–TCRβ+ T cells at days 0 and 5 from dissociated TA muscles of young, old, and geriatric mice (**Figure 2N**). We detected a low abundance of TCRβ+ T cells out of all CD45+ hematopoietic cells at day 0, but still detected a significantly higher T cell abundance in old mice compared to young (**Figure 2O**; Student’s t-test, p-value = 0.02*). Further, we observed an increase in T cell abundance from day 0 to day 5 and a significant difference between the old and young samples at 5 dpi (**Figure 2O**; Student’s t-test, p-value = 0.03*). Together, these results suggest the abundance of the T cell pool is elevated specifically in older muscles (20-months of age), but not preserved in geriatric ages.

Independent of age, we detected few MuSCs and progenitors immediately following injury (days 1 and 2; 2.1%, 1.1%, respectively) and the abundance of MuSCs and progenitors peaked at day 5 (9.7%) (**Supplementary Figure 4A**, **Supplementary Table 1A**). This was in agreement with previous studies.^20, 40^ The peak in MuSCs and progenitors abundance did vary by age with the young cells peaking at day 5, the old cells peaking at day 3.5, and the geriatric cells peaking at day 7 (**Figure 2I**). This difference in peak abundance did not result in a statistically significant difference in the MuSCs and progenitors dynamics by age, but it did demonstrate a delayed response by the geriatric MuSCs and progenitors. Independent of age, we detected the most Myonuclei at days 0 and 7 (15.1%, 10.6%, respectively) (**Supplementary Figure 4A**, **Supplementary Table 1A**). The Myonuclei dynamics were very similar between the three ages, but we detected more Myonuclei in old and geriatric samples at both days 0 and 7 compared to young (**Figure 2J**, **Supplementary Figure 5M**).

### Senescence scoring based on single-cell transcriptomic signatures

Next, we sought to investigate age-specific differences in senescence within skeletal muscle regeneration. Hallmarks of mammalian aging include stem cell exhaustion, altered cellular communication, and cellular senescence.^41^ Identifying senescent cells in scRNA-seq data is challenging because the markers traditionally used to identify senescent cells are either lowly expressed, expressed in select cell types in single-cell data, and/or assayed in terms of cellular localization and enzymatic function (**Figure 3A**).^42^ For example, senescent cells are commonly identified by persistent expression of cell cycle regulators p16 (*Cdkn2a*), p21 (*Cdkn1a*), p53, and/or Rb.^15^ Senescent cells are also marked by the senescent associated secretory phenotype (SASP) which includes proinflammatory cytokines and chemokines, growth modulators, angiogenic factors, and matrix metalloproteinases (e.g., *Mmp3*).^43^ To examine individual gene signatures of senescence, we quantified the abundance of *Cdkn2a* (encodes p16), *Cdkn1a* (encodes p21), *Mmp3* (a senescence-associated matrix metalloproteinase), *Glb1* (encodes senescence-associated ý-galactosidase) across all cell types and ages.

**Figure 3:**
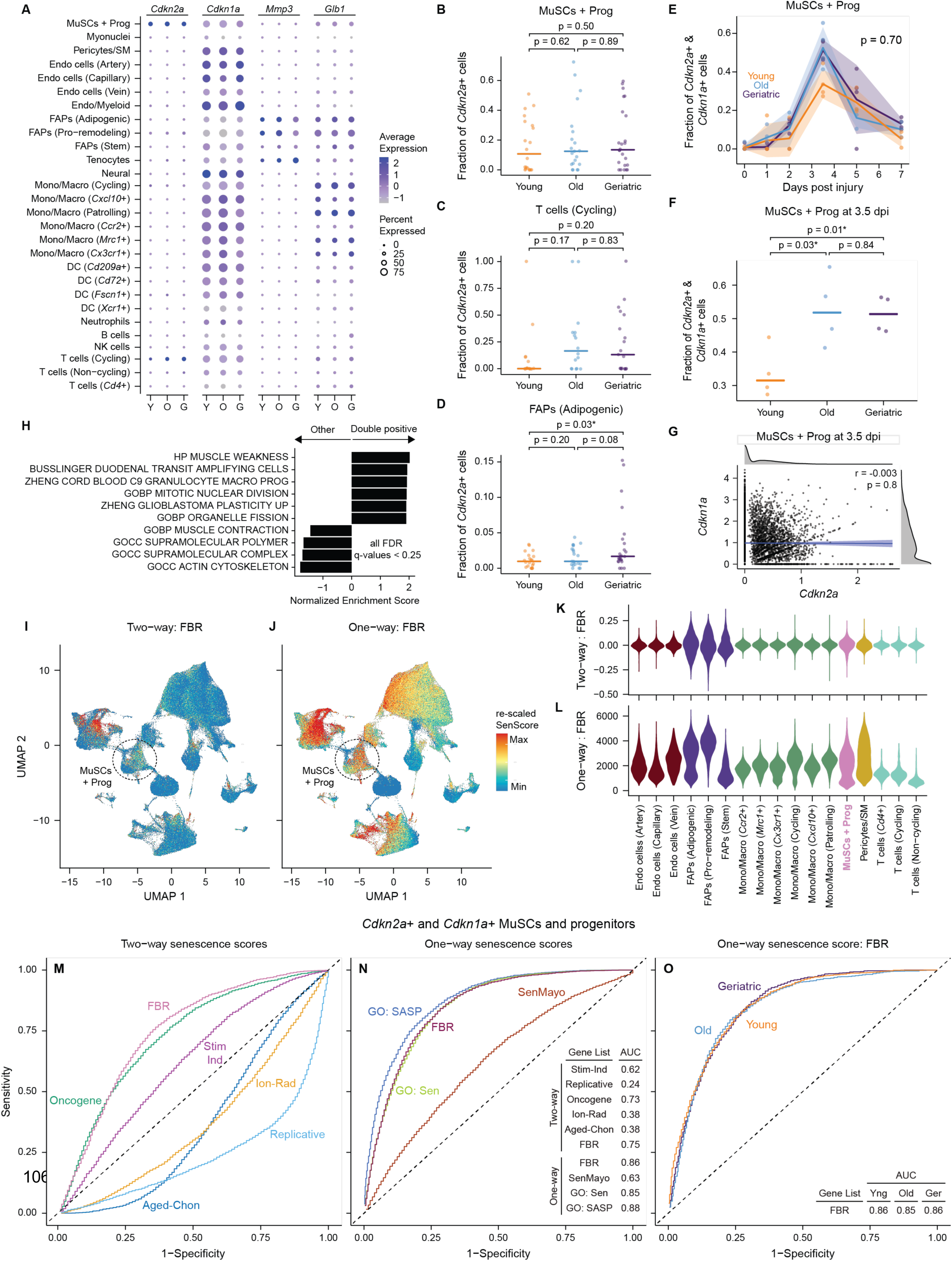
Identification of senescent-like cells using an informatic scoring approach. (**A**) Dot plot showing the expression frequency and average expression level of select senescence-associated genes by cell type and age (across all time-points). (**B**-**D**) Scatter plots of the fraction of *Cdkn2a*+ cells within the MuSCs and progenitors (**B**), T cells (Cycling; *Cd3e*+) (**C**), and FAPs (Adipogenic) (**D**) across all samples within each age group (n = 20-24). Points are the fraction for each sample. Horizontal line is the median for each age group. Significance was evaluated using the Student’s t-test. (**E**) Line plot of the fraction of MuSCs and progenitors that co-express *Cdkn2a* and *Cdkn1a* from 0-7 days post injury (dpi). Points are the fraction for each sample (n = 4). Ribbon is the standard deviation. Statistical significance of age-specific dynamics was evaluated using non-linear modeling and the FDR-corrected p-values is reported. (**F**) Scatter plot of the fraction of MuSCs and progenitors at 3.5 dpi that co-express *Cdkn2a* and *Cdkn1a* by age. Points are the fraction for each sample (n = 4). Horizontal line is the median for each age group. Significance was evaluated using the Student’s t-test. (**G**) Scatter plot of the normalized expression level of *Cdkn2a* and *Cdkn1a* transcripts in all individual MuSCs and progenitors at 3.5 dpi. The density is shown on the top and to the right of the plot. Blue line represents the linear trend. Ribbon is the confidence interval. The inset contains the Pearson correlation coefficient and its statistical significance. (**H**) Significantly up-or down-regulated gene ontology terms between *Cdkn2a*+ and *Cdkn1a*+ (Double Positive) and all other (Other) MuSCs and progenitors at day 3.5. The normalized enrichment score and the FDR-corrected q-values were obtained from gene set enrichment analysis (GSEA). (**I**-**J**) Feature plots of the final dataset with the cells colored by the re-scaled Two-way FBR senescence score (**I**) and the re-scaled One-way FBR senescence score (**J**). The cells are randomly plotted. (**K-L**) Violin plots of the Two-way (**K**) and One-way (**L**) FBR senescence scores in select cell types. (**M-O**) Receiver Operator Characteristic (ROC) curves based on the co-expression of *Cdkn2a* and *Cdkn1a* for the six Two-way senescence scores (**M**), the four One-way senescence scores (**N**), and for each age group using the One-way FBR senescence score (**O**). The area under the curve (AUC) is reported for each ROC curve.

Expression of these genes depended more on cell type than age, when considering all time-points and samples together (**Figure 3A**). Given *Cdkn1a*, *Mmp3*, and *Glb1* were widely expressed across many cell types, we focused on the common senescence hallmark gene *Cdkn2a*. *Cdkn2a* transcripts were rarely detected, in agreement with previous observations in the Tabular Muris Senis project^17^, and primarily observed in the MuSCs/progenitors, cycling T cells, and FAPs. We observed age-associated changes in the relative abundance of *Cdkn2a*+ cells within any given cell type infrequently significant (**Figure 3B-D**). There was no significant difference in the fraction of *Cdkn2a*+ MuSCs and progenitors by age or in the fraction of *Cdkn2a*+ cycling T cells by age (**Figure 3B-C**). However, we did observe a significant difference in the fraction of *Cdkn2a*+ adipogenic FAPs between the young and geriatric ages (**Figure 3D**; Student’s t-test, p-value = 0.03*).

Within the MuSCs and progenitors, we observed an increase in the fraction of cells that co-expressed *Cdkn2a* and *Cdkn1a* from day 0 to day 3.5, after which it returned to near day 0 levels (**Figure 3E**). Although there was no significant difference by age in the abundance of these double-positive cells across all timepoints, there was a significant difference at day 3.5 between the young and old ages (Student’s t-test, p-value = 0.03*) and the young and geriatric ages (Student’s t-test, p-value = 0.01*) (**Figure 3E-F**). Although *Cdkn2a* and *Cdkn1a* are both cell cycle inhibitors and senescent markers, their expression was not correlated in MuSCs and progenitors at day 3.5 on the individual cell level in this dataset, possibly due to transcript detection dropout (**Figure 3G**). We considered the double-positive *Cdkn2a+* and *Cdkn1a+* cells as candidate senescent MuSCs/progenitors and performed Gene Set Enrichment Analysis (GSEA) on day 3.5 at their peak abundance. GSEA found that double-positive MuSCs/progenitors are enriched for gene-sets associated with muscle weakness and various mitosis-related processes but diminished in muscle contraction and cytoskeletal processes (**Figure 2H**). Collectively, these GSEA results suggest that double-positive MuSCs/progenitors have signatures of dysregulated muscle function and stalled cell cycle-related gene expression.

We then used the *Cdkn2a*+ *Cdkn1a+* MuSCs/progenitors as a candidate cell population to evaluate broader senescence signatures at the single-cell level. We tested two senescence scoring methods^44, 45^ and ten senescence-signature (SenSig) gene lists^44, 46–49^ (**Extended Data File 2**). We refer to the first method as the Two-way Senescence Score (Sen Score) because it calculates a score based on a list of up- and down-regulated genes. Within this method we tested six gene lists that were generated from bulk RNA-seq datasets comparing cells with senescence conditions or markers to control cells from various tissues and cell types.^44, 46^ We refer to the second method as the One-way Sen Score because it calculates a score based on a list of up-regulated genes. Within this method we tested four gene lists, three of which were taken from gene-ontology databases or curated in other reports.^47–49^ We refer to the Methods**, Supplementary Figure 7**, and **Extended Data File 2** for more details on these two methods and the gene lists. We note that two of these gene lists are derived from bulk RNA-seq differential expression analyses of p16+ and p16– cells selected based on transgenic reporter status. One gene list (“FBR”) was generated by Cherry et al from p16+ versus p16– CD29+ cells isolated from a foreign body response-driven skeletal muscle fibrosis model in adult *p16-CreER^T^*^2^*;Ai14* reporter mice.^44^ A second gene list (“Aged Chondrocytes”) was generated from p16+ versus p16– Aggrecan+ chondrocytes isolated from 20-mo *p16-tdTom;Aggrecan-CreER^T^*^2^*;Ai6* mice (B.O. Diekman, personal communication).^50–52^ To compare how the choice of gene list and method impacted senescence scoring across different cell types, we examined the FBR two-way and one-way scores in the final scRNA-seq dataset. The Two-way FBR scores were more consistently low across most cell clusters and exhibited high scores most notably in the FAP and MuSC clusters (**Figure 3I,K**). The One-way FBR scores had broader distribution, with many more cell types exhibiting high scores, including FAP, MuSCs, endothelial cell, pericytes and smooth muscle cell, and monocytes/macrophages clusters (**Figure 3J,L**). Whereas the Two-way FBR scores were mean-centered around zero from each cell type cluster due to their z-scored counts (**Figure 3K**), One-way FBR scores had varied cell type averages (**Figure 3L**). These differences complicate establishing threshold for senescence positivity between cell types in the one-way scores.

We established a scoring approach calibrated for sensitivity and specificity in discriminating *Cdkn2a*+ *Cdkn1a+* MuSCs/progenitors across all ages and timepoints and present the results in receiver-operator curves with performance reported using an area-under-the-curve (AUC) metric (**Figure 3M-O**). Between the two senescence scoring methods and the ten SenSig gene sets, the One-way FBR method performed the best of any experimentally derived approach (AUC = 0.86), and was comparable to the ontology-curated One-way GO: SASP approach (AUC = 0.88; **Figure 3M-N**). Notably, it performed far better than the recently described SenMayo list while using the same ssGSEA method (AUC = 0.63). Moreover, the One-way FBR score accurately discriminated double-positive senescent-like MuSCs across all three ages (AUCs = 0.85-0.88), suggesting it captures common features of senescence irrespective of age (**Figure 3O**). We concluded that the One-way FBR method was able to accurately identify senescent-like cells in a manner that is not biased by a highly curated gene list.

### Refined analysis of myogenic subsets

In the final dataset, we identified two broad myogenic clusters (**Figure 1F**). We observed a cluster of MuSCs and progenitors that expressed the myogenic transcription factor *Pax7*^53^ and a cluster of myonuclei that expressed *Acta1*, *Myh1*, and *Myh4*, genes critical for the contractile function of mature skeletal muscle cells^54^ (**Supplementary Figure 3C**). We subsetted out these myogenic clusters and re-clustered and re-embedded the cells, resulting in nine distinct sub-clusters for refined annotation (**Figure 4A-B**). We identified four progenitor populations that expressed *Pax7*, *Myf5*, *Myod1*, *Myog*, *Mymk*, and *Mymx* and three myonuclei subtypes (IIx, IIb, IIx/IIb) that expressed *Acta1*, *Ckm*, and *Tnnt3*.^54^ We observed two transcriptomically variant clusters, which expressed both myogenic markers and either endothelial cell markers like *Cd34*, *Cdh5*, and *Pecam1* or monocyte and macrophage markers like *Ccr2* and *C1qa* (**Supplementary Figure 8**). We suspected these clusters were dominated by doublets. The first doublet sub-cluster contained cells that co-expressed *Pax7* and *Pecam1*, suggesting that these were MuSCs/progenitors and endothelial cell doublets (**Supplementary Figure 8A**). The second doublet sub-cluster contained cells that co-expressed *Acta1* and *C1qa*, suggesting that these were myonuclei and monocyte/macrophage doublets (**Supplementary Figure 8B**). We designated these sub-clusters as ‘Doublets 1’ and ‘Doublets 2’, respectively, and excluded them from subsequent analyses involving the myogenic subset.

**Figure 4:**
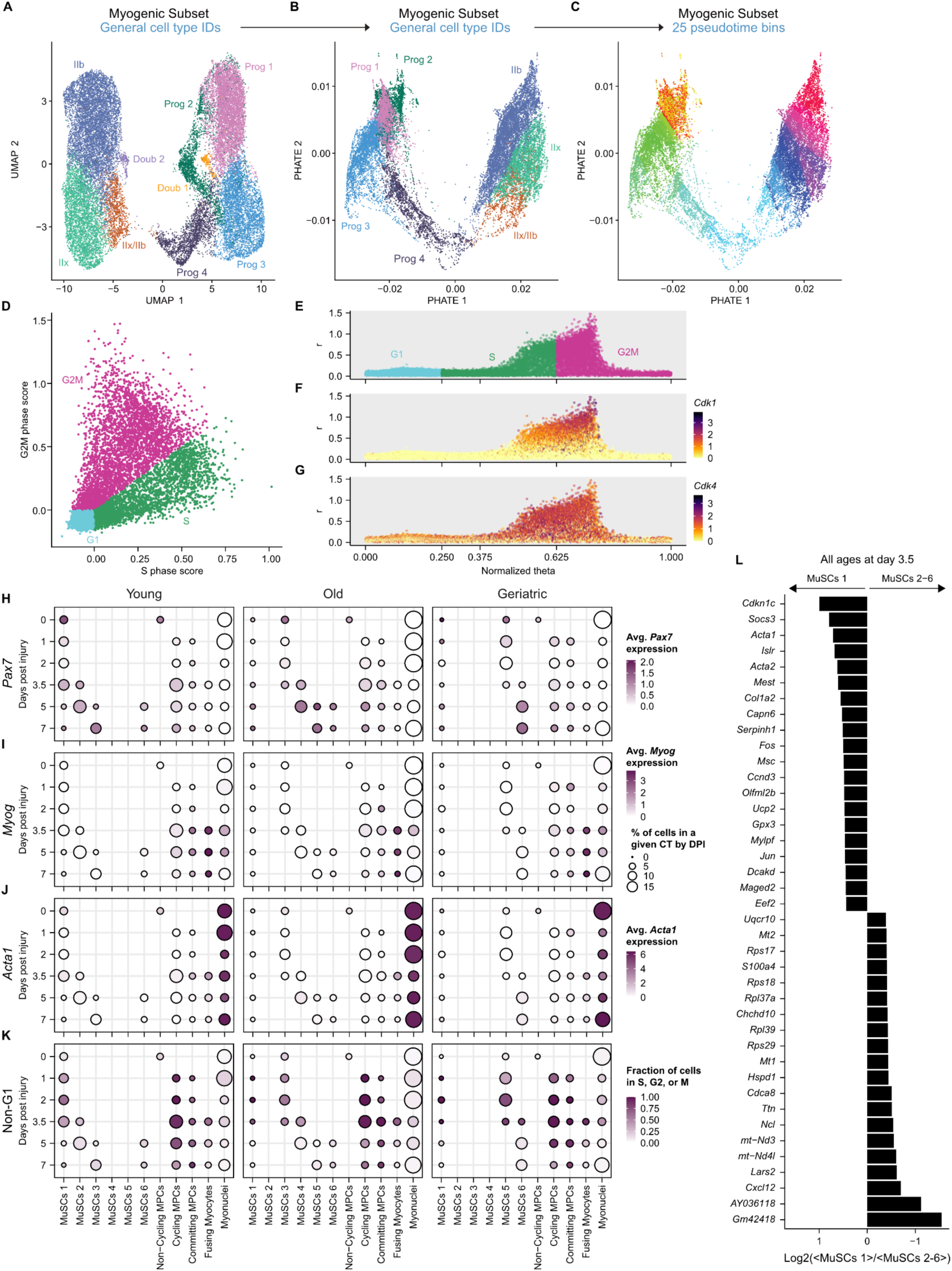
Age-specific trajectories through myogenesis following injury. (**A**-**C**) Pseudotime organization of the myogenic cell subset after re-clustering. Myogenic cells were re-embedded with UMAP (**A**) and with PHATE (**B**) and are colored by general myogenic IDs. PHATE embeddings were used by Monocle3 to organize the cells in pseudotime where the earliest pseudotime is in the upper left corner and the latest pseudotime in the upper right corner. Myogenic cells were organized into 25 approximately equal bins of increasing pseudotime values (**C**). (**D**) Each cell was assigned an S phase score and a G2M phase score using Seurat’s Cell Cycle Scoring method. Scatter plot of the two scores with the cells colored by the predicted cell cycle phase based on the two scores. (**E**-**G**) Polar coordinates (in **D**) were converted to cartesian coordinates and rescaled to fit in a range from 0 to 1. Cells are colored by the predicted cell cycle phase based on the two scores (**E**) and by the log-normalized expression of *Cdk1* (**F**) and *Cdk4* (**G**). (**H**-**K**) Dot plots of the average log-normalized expression of *Pax7* (**H**), *Myod1* (**I**), and *Acta1* (**J**) and by the fraction of non-G1 (S, G2, and M) cells (**K**) in each day post injury (dpi) and pseudotime-based myogenic cell state bin. The size of the circle is the percent of cells in each pseudotime-based myogenic cell state bin for each age and dpi combination. (**L**) Top 40 differentilally up- and down-regulated genes between cells in MuSCs 1 and MuSCs 2-6 at day 3.5. All genes highlighted here have an FDR-corrected q-value < 0.05.

The type IIx cluster expressed *Myh1*, the type IIb cluster expressed *Myh4*, and the type IIx/IIb cluster expressed both *Myh1* and *Myh4* (**Figure 4B**, **Supplementary Figure 9A-G**).^55^ A previous study also identified a myonuclei cluster that expressed both *Myh1* and *Myh4*, but this studied concluded that this cluster represented cells with high metabolic activity, not cells that represent a transitional state between types IIx and IIb.^18^ We interpret that the type IIx/IIb cluster identified here represents a transitional state between types IIx and IIb because 39% of the cells in this cluster co-express *Myh1* and *Myh4* and because this cluster does not differentially express markers of high metabolic activity (*Tnnc2*, *Tnni2*, *Mb*, *Cox6a2*, *Cox6c*, *Atp5e*, *Atp5g1*) (**Supplementary Figure 9F-H**).^18, 56, 57^ Additionally, this transitional fiber type is the most common transitional fiber type in rat and mouse muscle fibers.^58, 59^ Notably, the type IIx/IIb cluster had a lower percentage of mitochondrial reads than the type IIx and type IIb clusters, indicating that these cells were not clustering together due to being lower quality (**Supplementary Figure 9I**). We did not identify a cluster of neuro-muscular junction or myotendinous junction cells due to a lack of *Chrne* or *Col22a1* expression (**Supplementary Figure 9H**).^18, 29^

We then re-embedded the myogenic subset using PHATE^60^ and these embeddings were used by Monocle3^61–63^ to organize the cells in pseudotime (**Figure 4B-C**). The pseudotime values were grouped into 25 bins that contain approximately equal numbers of cells. As the cells progressed through pseudotime, the order in which myogenic markers were expressed followed a typical trajectory of myogenesis. Early pseudotime bins had predominant expression of *Pax7* and *Myf5*, but no strong expression of activation markers (**Supplementary Figure 10A-F**). Further in the pseudotime progression, cells still expressed *Pax7* and *Myf5*, but they also expressed cycling markers such as *Cdk1/4*. In later pseudotime bins, cells expressed *Myod1*, *Myog*, *Mymx*, and *Mymk*, markers of committed and fusing progenitors. In the latest pseudotime bins, cells expressed *Acta1*, *Ckm*, *Myh1*, and *Myh4*, markers of myogenic maturation.

### Pseudo-temporal analysis of myogenesis progression across mouse age and injury time

To directly compare myogenesis in regeneration responses, we assembled an annotated “cartography” of myogenic progression arrayed across both day post-injury and myogenic pseudotime. We first examined the percent of cells that fell within each of the 25 initial pseudotime bins by time-point and age, and then used expression frequency of myogenic marker genes to inform myogenic cell-state annotations (**Supplementary Figure 10A-H**). Pseudotime bins 1-6 exhibited age-specific cell abundances, with bins 1-2 predominantly containing cells from young mice, bin 3 from young and old mice, bin 4 contained cells from old mice, bin 5 from old and geriatric mice, and bin 6 from geriatric mice only. Bins 1-6 cells expressed *Pax7* and *Myf5*, which we annotated as MuSCs (with sub-stages 1-6 preserved). Bins 7-13 at 0 dpi expressed *Pax7*, *Myf5*, and *Myod1* and lowly expressed cycling markers *Cdk1* and *Cdk4*, which we annotated as Non-cycling MPCs. Cells in bin 7 and dpi 1-3.5 and cells in bins 8-11 and dpi 1-7 expressed *Myod1* and cycling markers *Cdk1* and *Cdk4*, which we annotated as Cycling MPCs. Cells in bin 12 at dpi 0-7 and cells in bin 13 at dpi 1-2 have diminishing expression of *Myf5*, *Cdk1* and *Cdk4*, and increasing expression of *Myog* and *Mymk*, which we annotated as Committing MPCs. Cells in pseudotime bin 13 and dpi 3.5-7 highly expressed *Myog* and *Mymk* and lowly expressed *Cdk1* and *Cdk4*, which we annotated as Fusing Myocytes. Cells in pseudotime bins 14-25 and dpi 0-7 expressed *Acta1*, which we annotated as Myonuclei. These pseudotime-informed myogenic cell stage aggregates (summarized in **Supplementary Figure 10I**) were used in subsequent analyses.

To infer the cell-cycle phases, we first assigned each myogenic cell S-phase and G2M-phase scores using Seurat’s standard Cell-Cycling Scoring method^64^ (**Figure 4D**). We treated these scores as polar coordinates, which were converted to cartesian coordinates and normalized to be within a range from 0 to 1 (“normalized theta”; **Figure 4E-G**, **Supplementary Figure 11A-B**). We assessed all myogenic cells within this cell cycle progression from a normalized theta of 0 to 1, corresponding to the continuum of G1–S–G2M stages (**Figure 4E**, **Supplementary Figure 11A-B**). When considering all MuSCs/progenitors, the distribution of normalized theta values increased from day 0 to day 3.5 and nearly returned to day 0 levels by day 7 in all age groups, suggesting a return to quiescence as expected (**Supplementary Figure 11E**). We observed a shift to higher normalized theta values at 1 dpi in the geriatric samples compared to the young and old myogenic cells, suggesting an age-skewed cell-cycle induction in early injury-response that may represent a precocious activation phenotype. Differences by age group were minimal after 3.5 dpi. When examining all myogenic cells by the 25 pseudotime bins, we observed a shift in normalized theta values at bin 7 persisting through bin 13 (**Supplementary Figure 11F**). Notably, pseudotime bins 7-13 also highly expressed the cycling markers *Cdk1* and *Cdk4* (Supplementary Figure 10G-H).

We then found that *Cdk1* and *Cdk4* are more highly expressed in cells predicted to be in S or G2M than in cells predicted to be in G1 (**Figure 4F-G**). Seurat’s standard G1 cutoff is at a normalized theta value of 0.25. Based on the expression of cycling markers *Cdk1* and *Cdk4* and the distribution of cells across the normalized theta values, we extended the G1 cutoff to 0.375 for this analysis (**Figure 4F-G**, **Supplementary Figure 11C-D**). For simplicity, cells with a normalized theta below and above 0.375 were classified as “G1” and “Non-G1” (S/G2/M), respectively (**Supplementary Figure 11C-D**).

We then calculated the percent of cells within each myogenic cell stage and time-point by age group. Within the MuSC 1-6 stages at 0-1 and 5-7 dpi, we observed high levels of *Pax7* (**Figure 4H**). We also detected a high fraction of Non-G1 cells in MuSCs 1-6, especially at 1-3.5 dpi in the old and geriatric mice (**Figure 4K**). This suggested that more of the old and geriatric MuSCs were actively cycling post-injury compared to the young MuSCs. We compared the 3.5 dpi cells in the MuSC 1 and MuSC 2-6 stages by differential gene expression and found that the quiescence-associated genes *Cdkn1c* (encoding p57^Kip2^) and *Socs3* were upregulated in MuSCs 1 and numerous translation-associated genes such as *Rps29* were upregulated in MuSCs 2-6 (**Figure 4L**). These expression profiles suggest that MuSCs 1 cells are in a less activated state MuSCs 2-6 cells.

We observed an inverse relationship between the average *Myog* expression and the fraction of Non-G1 cells in the Cycling MPCs, Committing MPCs, and Fusing Myocytes, as expected for differentiating myogenic cells (**Figure 4I,K**). In all ages we detected the highest average expression of *Myog* in the Fusing Myocytes population (**Figure 4I**). We detected Myonuclei in all ages and at every dpi, but there were fewer Myonuclei with lower average expression of *Acta1* at dpi 1-3.5 in the geriatric mice compared to the young and old mice (**Figure 4J**). Together, these results present an integrated cellular cartography of myogenic trajectories through regeneration, which exhibits age-associated cellular trajectories, particularly within the MuSC pool.

### Scoring cell senescence across the myogenic cell cycle

To explore how cellular senescence manifests within this organized cartography of myogenesis, we focused on the One-way FBR Sen Score which performed well across ages (**Figure 3O**). To identify cells with senescence-like identities, we defined a threshold within the One-way FBR Sen Score based on its relationship with *Cdkn2a* and *Cdkn1a* expression which exhibited correlation (**Supplementary Figure 11G-H**). We set a One-way FBR score threshold at 2412, where 50% of cells above this value co-expressed *Cdkn2a* and *Cdkn1a,* and classified cells above this threshold as ‘Sen Score high’ and senescence-like (**Figure 5A**). We further observed through ROC analysis that the One-way FBR Sen Score could accurately identify double-positive *Cdkn2a* and *Cdkn1a* cells from both the G1 and Non-G1 fractions of MuSCs/progenitors (**Figure 5B**).

**Figure 5:**
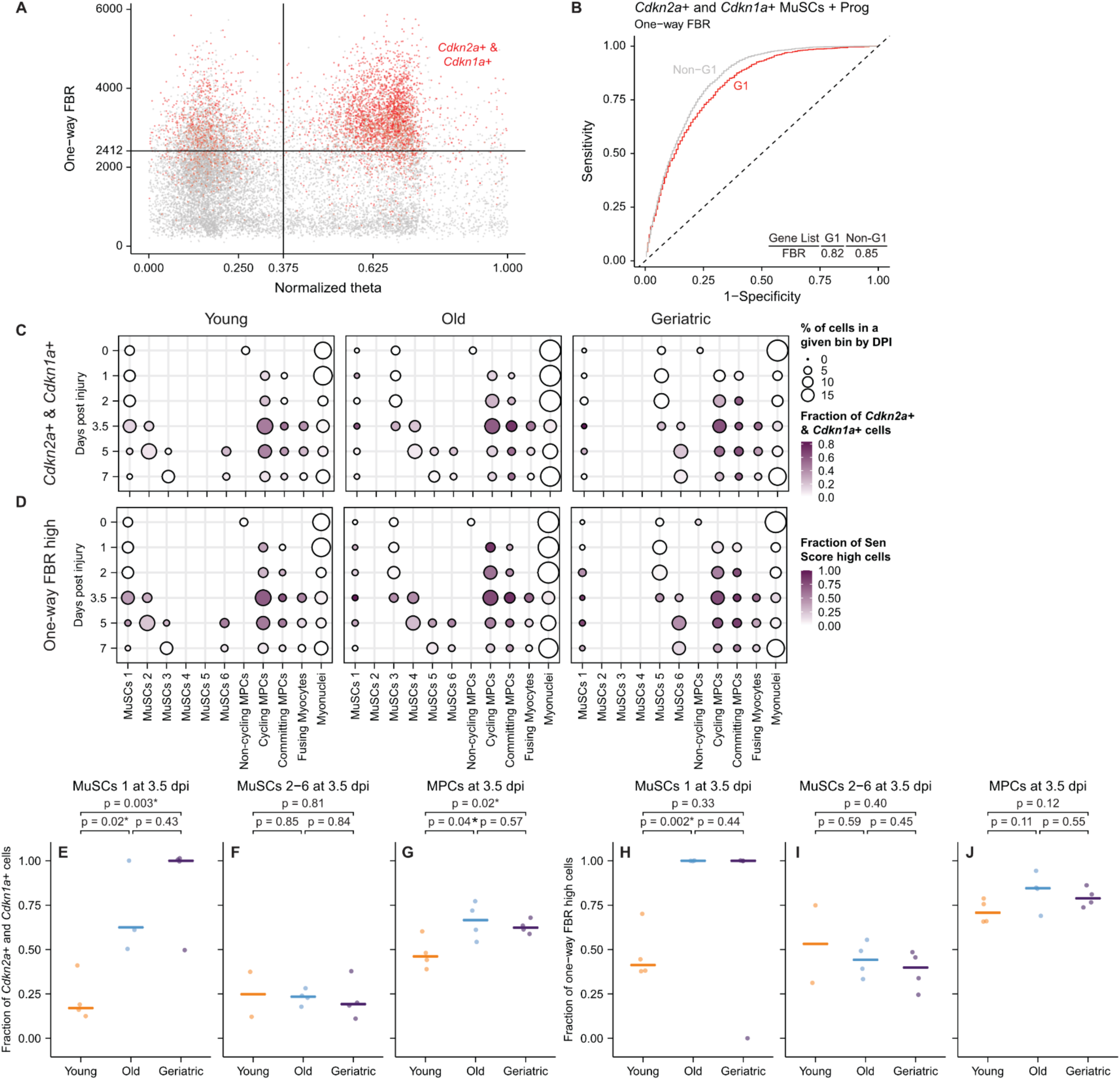
Aging-associated accumulation of senescent-like cells at critical transitory myogenic cell states. (**A**) Scatter plot of the normalized cartesian coordinate Cell Cycle Scores and the One-way FBR senescence score in all MuSCs and progenitors. Cells colored in red co-express *Cdkn2a* and *Cdkn1a*. All other cells in gray. The vertical line is the G1 cutoff, and the horizontal line is where 50% of the cells above this line co-express *Cdkn2a* and *Cdkn1a*. (**B**) Receiver Operator Characteristic (ROC) curves based on the co-expression of *Cdkn2a* and *Cdkn1a* in the MuSCs and progenitors (all ages and days post-injury (dpi)) in G1 and non-G1 (S, G2, and M) when using the One-way FBR senescence score. Area under the curve is reported for each ROC curve. (**C**-**D**) Dot plots of the fraction of cells that co-express *Cdkn2a* and *Cdkn1a* (**C**) and the fraction of One-way FBR senescence score-high cells (**D**) in each dpi and pseudotime-based myogenic cell state bin. The size of the circle is the percent of cells in each pseudotime-based myogenic cell state bin for each age and dpi combination. (**E**-**G**) Scatter plot of the fraction of MuSCs 1 (**E**), MuSCs 2-6 (**F**), and all MPCs (**G**) at 3.5 dpi that co-express *Cdkn2a* and *Cdkn1a* by age. Points are the fraction for each sample (n = 2-4) and the horizontal line is the median for each age group. Significance was evaluated using the Student’s t-test. (**H**-**J**) Scatter plot of the fraction of MuSCs 1 (**H**), MuSCs 2-6 (**I**), and all MPCs (**J**) at 3.5 dpi that are One-way FBR senescence score high by age. Points are the fraction for each sample (n = 2-4) and the horizontal line is the median for each age group. Significance was evaluated using the Student’s t-test.

We aimed to quantify the prevalence of cellular senescence within the cartography of myogenesis by age using both double-positive expression status and Sen Score, reasoning that the scores might capture a more expansive set of senescence-like cells. We observed a correspondence between the fraction of cells that co-express *Cdkn2a* and *Cdkn1a* and the fraction of cells that have a high Sen Score in most ages, cell stages, and timepoints (**Figure 5C-D**). Focusing on 3.5 dpi, we found a significantly higher fraction of *Cdkn2a*+ and *Cdkn1a*+ cells in both the old and geriatric MuSCs 1 compared to young MuSCs 1 (Student’s t-tests, p-values = 0.02* and 0.003*, respectively; **Figure 5E**). We did not observe age-specific differences in the fraction of *Cdkn2a*+ and *Cdkn1a*+ in MuSCs 2-6 (**Figure 5F**). We found a significantly higher fraction of *Cdkn2a*+ and *Cdkn1a*+ in both old and geriatric MPCs (from both Cycling and Committing stages) compared to young MPCs (Student’s t-tests, p-values = 0.04* and 0.02*, respectively; **Figure 5G**). Likewise, we observed a similar pattern in the fraction of One-way FBR Sen Score high cells, with significantly higher frequencies in the old compared to young MuSCs 1 (Student’s t-test, p-value = 0.002*; **Figure 5H**). We did not observe age-specific differences in the fraction of Sen Score high MuSCs 2-6 or MPCs (**Figure 5I-J**). Together, these observations point to a transitory senescent-like cell population that is abundant at the self-renewing MuSC 1 stage across all ages, but increases in older mice, potentially underlying a stalled stem-cell self-renewal in mouse muscle aging.

## DISCUSSION

We utilized the profiling depth and complexity of scRNA-seq and associated computational analyses to generate a comprehensive compendium of 273,923 single-cell transcriptomes from regenerating tibialis anterior muscles throughout mouse lifespan. To date, our dataset is the most comprehensive portrait of muscle repair at the single-cell level, as it includes three ages (young, old, and geriatric), six time points (days 0, 1, 2, 3.5, 5, and 7), and includes 29 different cell type clusters (**Figure 1**, **Supplementary Figures 1** and **3**). Additionally, compared to previous scRNA-seq and snRNA-seq skeletal muscle studies, we have identified more specific endothelial, FAPs, and immune cell sub-types.^11, 18–30^

The immune, stromal, and myogenic cells found in skeletal muscle contribute to muscle maintenance and regeneration by regulating MuSC quiescence, proliferation, and differentiation.^1^ It has been shown that an imbalance in immune cell populations during injury response can disrupt proper muscle repair.^1, 2^ To investigate this we compared the change in cell type abundances over our regeneration time course between young, old, and geriatric muscles. As expected, Neutrophils are one of the first immune cell types to peak in abundance (**Supplementary Figure 5L**).^4^ We also observe monocyte and macrophage populations that express pro-inflammatory markers like *Ccr2* and patrolling markers like *Ctsa* responding soon after injury (days 1-2) when we expect the muscle environment to be enriched with pro-inflammatory cytokines (**Figure 2A,E**).^1, 4^ Monocytes and macrophages that express pro-inflammatory markers clear cellular debris and promote myogenic cell proliferation.^1, 5^ There should be a shift to monocytes and macrophages that express anti-inflammatory marker *C1qa* at 4-7 dpi (**Figure 2B-C**, **Supplementary Figure 3A**).^4^ We do broadly observe a shift from monocytes and macrophages that express pro-inflammatory markers to anti-inflammatory markers, but there are significant differences by age (**Figure 2A-E**). This difference in monocyte and macrophage dynamics could explain the age-related decline in muscle repair because if macrophages do not clear cellular debris or promote myogenic cell proliferation and differentiation, the muscle remains inflamed and there are repeated cycles of necrosis and regeneration.^5^ The damaged myofibers are then replaced with adipose tissue, fibrotic tissue, or bone, instead of new myofibers.^5^

In addition to age-specific differences in the dynamics of the monocyte and macrophage populations, we observe age specific differences in the T cell dynamics (**Figure 2L-O**, **Supplementary Figure 5T**). It has previously been shown that Treg cells, marked by *Cd4* and *Foxp3*, accumulate in injured muscle and to peak in abundance at day 4.^4^ We detected *Foxp3* expression in a few T cells, specifically in the T cell (Cycling) and T cell (Non-Cycling) populations which both highly express *Cd3e* (**Supplementary Figure 3A**). Although we cannot confidently identify any of our three T cell populations as Tregs, we do observe a peak in T cell abundances at days 5 and 7 (age-specific). There is miscoordination of T cell response which in turn could impact the ability of aged muscle to repair itself.

One factor that has been shown to contribute to the reduced functionality of MuSCs in aged tissues is the establishment of senescent MuSCs.^2, 13^ Prior studies have used the cell cycle proteins p16, p21, p53, and Rb to differentiate between dividing and non-diving cells, but the non-diving cells can include senescent cells and quiescent cells.^15^ SA-β-gal is also commonly used to identify senescent cells, but it is also detected in quiescent cells and in stressed cells.^15, 65^ Because these markers are not unique to senescent cells and because senescent cells are heterogeneous, it has been challenging to identify biomarkers that can accurately and consistently identify senescent cells across species, tissues, and conditions.^16^ Indeed, recent large consortia have been established to develop new tools to detect and bioinformatically identify senescent cells with robustness and precision throughout mammalian tissues and lifespans.^66^ Given the role that cellular senescence plays in limiting cell contributions in aged tissues, we tested a series of tools to bioinformatically identify senescence in these single-cell data and found a transfer-learning based scoring approach accurately classified senescent-like myogenic cells across ages and cell cycling states. The approach described here to quantitatively assess various senescence scoring approaches and reference gene lists in discriminating senescent or senescent-like cells (**Figures 3** and **5**) may provide a template for future studies using single-cell data. Notably, here we concluded that a skeletal muscle FBR gene list more accurately and robustly discriminated senescent-like *Cdkn2a*+ and *Cdkn1a*+ MuSC/progenitors in these muscle regeneration datasets than did a variety of experimental and curated gene lists, including the recently described SenMayo list.^47^ In particular, at day 3.5 post-injury, we observed a significantly higher fraction of *Cdkn2a*+ and *Cdkn1a*+ cells in the aged and geriatric MuSCs associated with a self-renewing cell stage (MuSC stage 1; **Figure 5E,G**). Likewise, we observed a similar pattern in the fraction of One-way FBR Sen Score high cells, with significantly higher frequencies in the old compared to young MuSCs 1 (**Figure 5H**). These observations point to a transitory senescence-like cell population that is abundant at the self-renewing MuSC stage across all ages, but increases in older mice, potentially underlying a stalled stem-cell self-renewal in mouse muscle aging.

## METHODS

### Mouse muscle injury and single-cell isolation

Muscle injury was induced in young (4-7 months-old [mo]), old (20 mo), and geriatric (26 mo) C57BL/6J mice (Jackson Laboratory # 000664; NIA Aged Rodent Colonies) by injecting both tibialis anterior (TA) muscles with 10 µl of notexin (10 µg/ml; Latoxan, France). The mice were sacrificed, and TA muscles were collected at 0, 1, 2, 3.5, 5, and 7 days post-injury (dpi). Each TA was processed independently to generate single cell suspensions. At each time point, the young and old replicates are biological replicates, and the geriatric replicates are two pairs of technical replicates (n = 3-4). Muscles were digested with 8 mg/ml Collagenase D (Roche, Basel, Switzerland) and 10 U/ml Dispase II (Roche, Basel, Switzerland) and then manually dissociated to generate cell suspensions. Myofiber debris was removed by filtering the cell suspensions through a 100 µm and then a 40 µm filter (Corning Cellgro # 431752 and # 431750). After filtration, erythrocytes were removed by incubating the cell suspension in erythrocyte lysis buffer (IBI Scientific # 89135-030).

### Single-cell RNA-sequencing library preparation

After digestion, the single-cell suspensions were washed and resuspended in 0.04% BSA in PBS at a concentration of 10^6^ cells/ml. A hemocytometer was used to manually count the cells to determine the concentration of the suspension. Single-cell RNA-sequencing libraries were prepared using the Chromium Single Cell 3’ reagent kit v3 (10x Genomics, Pleasanton, CA) following the manufacturer’s protocol.^67^ Cells were diluted into the Chromium Single Cell A Chip to yield a recovery of 6,000 single-cell transcriptomes with <5% doublet rate. Libraries were sequenced on the NextSeq 500 (Illumina, San Diego, CA).^68^ The sequencing data was aligned to the mouse reference genome (mm10) using CellRanger v5.0.0 (10x Genomics).^67^

### Preprocessing and batch correction of single-cell RNA sequencing data

From the gene expression matrix, the downstream analysis was carried out in R (v3.6.1). First, ambient RNA signal was removed using the default SoupX (v1.4.5) workflow (autoEstCounts and adjustCounts; github.com/constantAmateur/SoupX).^36^ Samples were then preprocessed using the standard Seurat (v3.2.3) workflow (NormalizeData, ScaleData, FindVariableFeatures, RunPCA, FindNeighbors, FindClusters, and RunUMAP; github.com/satijalab/seurat).^64^ Cells with fewer than 200 genes, with fewer than 750 UMIs, and more than 25% of unique transcripts derived from mitochondrial genes were removed. After preprocessing, DoubletFinder (v2.0.3) was used to identify putative doublets in each dataset.^37^ The estimate doublet rate was 5% according to the 10x Chromium handbook. The putative doublets were removed from each dataset. Next, the datasets were merged and then batch-corrected with Harmony (github.com/immunogenomics/harmony) (v1.0).^38^ Seurat was then used to process the integrated data. Dimensions accounting for 95% of the total variance were used to generate SNN graphs (FindNeighbors) and SNN clustering was performed (FindClusters). A clustering resolution of 0.8 was used resulting in 24 initial clusters.

### Cell type annotation in single-cell RNA sequencing data

Cell types were determined by expression of canonical genes. Each of the 24 initial clusters received a unique cell type annotation. The nine myeloid clusters were challenging to differentiate between, so these clusters were subset out (Subset) and re-clustered using a resolution of 0.5 (FindNeighbors, FindClusters) resulting in 15 initial clusters. More specific myeloid cell type annotations were assigned based on expression of canonical myeloid genes. This did not help to clarify the monocyte and macrophage annotations, but it did help to identify more specific dendritic cell and T cell subtypes. These more specific annotations were transferred from the myeloid subset back to the complete integrated object based on the cell barcode.

### Analysis of cell type dynamics

We generated a table with the number of cells from each sample (n = 65) in each cell type annotation (n = 29). We removed the erythrocytes from this analysis because they are not a native cell type in skeletal muscle. Next, for each sample, we calculated the percent of cells in each cell type annotation. The mean and standard deviation were calculated from each age and time point for every cell type. The solid line is the mean percentage of the given cell type, the ribbon is the standard deviation around the mean, and the points are the values from individual replicates. We evaluated whether there was a significant difference in the cell type dynamics over all six time points using non-linear modeling. The dynamics for each cell type were fit to some non-linear equation (e.g., quadratic, cubic, quartic) independent and dependent on age. The type of equation used for each cell type was selected based on the confidence interval and significance (p < 0.05) for the leading coefficient. If the leading coefficient was significantly different from zero, it was concluded that the leading coefficient was needed. If the leading coefficient was not significantly different than zero, it was concluded that the leading coefficient was not needed, and the degree of the equation went down one. No modeling equation went below the second degree. The null hypothesis predicted that the coefficients of the non-linear equation were the same across the age groups while the alternative hypothesis predicted that the coefficients of the non-linear equation were different across the age groups. We conducted a likelihood ratio test to see if the alternative hypothesis fits the data significantly better than the null hypothesis and we used FDR as the multiple comparison test correction.

### Muscle immunohistochemical analysis

Muscle injury was induced in young, old, and geriatric C57BL/6J mice by injecting both TA muscles with 10 µl of notexin (10 µg/ml; Latoxan, France). The mice were sacrificed, and TA muscles were collected at 0 and 5 dpi. The TA muscles were coated in Tissue-Tek O.C.T. Compound (Sakura Finetek # 4583), snap-frozen in liquid nitrogen cooled isopentane (Thermo Scientific Chemicals # AA19387AP), and then stored at -80C. Frozen TA muscles were sectioned with a cryostat transversely at 5 µm thickness and section slides were stored at –20C until stained. Sections were fixed with 4% PFA (Electron Microscopy Sciences # 15710) for 10 minutes, washed with 1X PBS, and blocked with 3% BSA (Rockland Immunochemicals # RLBSA50) at room temperature for 1 hour. Sections were washed with 1X PBS and then stained with rat anti-mouse CD3 (eBiosciences # 14-0032-82) at 1:100 dilution in blocking buffer overnight at 4C. Sections were then washed and stained with Alexa Fluor Plus 750 Phalloidin (Life Technologies # A30105) at 1:500 dilution and goat anti-rat 488 (Invitrogen # A-11006) at 1:250 dilution in blocking buffer for 1 hour at room temperature protected from light. Sections were then washed with 1X PBS and stained with 5 mg/mL DAPI (Life Technologies # D3571) at 1:1000 dilution. Slides were mounted with Glycergel mounting medium (Agilent # C056330-2) and stored at 4C before imaging. Images were acquired using a Nikon Eclipse Ti-E microscope (Micro-Video Instruments, Inc.), and were analyzed using NIS Elements 5.11.03 software and ImageJ 2.1.0.

### Immune cell flow cytometric analysis

Single cell suspensions of uninjured day 0 and injured day 5 TAs and gastrocnemius muscles were collected in the same way as the single cell suspensions for scRNA-seq library preparation. However, the single cells were suspended in 90% FBS and 10% DMSO and frozen at –80C. When ready to use, the single cell suspensions were thawed in a 37C water bath and then transferred to 15 mL conical tubes. The cells were washed with staining buffer (1X PBS + 0.5% BSA + 2 mM EDTA) before being spun at 500g for three minutes and transferred to a 96 well round bottom plate. Cells were incubated with FC block (TruStain FcX PLUS (anti-mouse CD16/32), Biolegend # 156604) at 4C for five minutes. Cells were then washed with staining buffer, spun at 500g for three minutes, and incubated with viability dye (Fixable viability dye APCe780, eBiosciences # 65-0865-14) and surface antibody (see antibody table) in BSB (Brilliant Stain Buffer Plus, BD Biosciences # 566385) and staining buffer for 30 minutes at 4C in the dark. Aliquots of cell samples were counted on a MoxiZ Mini Automated Cell Counter. After incubating we followed the manufacturer’s protocol (FOXP3 Transcription factor fixation/permeabilization kit, eBioscience # 00-5521-00) and cells were washed with staining buffer, spun, and resuspended in FoxP3 1x perm solution (10x Permeabilization buffer, Invitrogen # 00-8333-56) and incubated for 30 minutes at 4C in the dark. Cells were washed with 1x perm solution, spun twice, and resuspended in an intracellular antibody stain. Cells were incubated in intracellular antibody stain for 30 minutes at 4C in the dark. Cells were washed with 1x perm and spun twice before being resuspended in staining buffer and transferred to 40 µM blue capped flow tubes. FMO controls were prepared using a mixture of young, old, and geriatric uninjured and injured TA and gastrocnemius muscles. Flow cytometry was performed using a FACSymphony A3 (BD) and data was analyzed in FlowJo 10.5.3. Gates were determined using FMO controls.

### Flow cytometry antibodies

| Antibody | Clone | Fluorophore | Manufacturer catalog |
| --- | --- | --- | --- |
| CD45 | 30-F11 | PECy7 | Invitrogen # 25-0451-82 |
| CD11b | M1/70 | BUV395 | BD Biosciences # 563553 |
| CD11c | N418 | PerCPCy5.5 | Invitrogen # 45-0114-82 |
| TCRβ | H57-597 | APC | eBiosciences # 17-5961-82 |
| CD19 | 1D3 | BV650 | BD Biosciences # 563235 |

#### Senescence scoring

We tested two senescence scoring methods along with ten senescence gene lists (Extended Data File 2) to identify senescent-like cells within the scRNA-seq dataset. The Two-way senescence score was calculated using a transfer-learning method developed by Cherry et al 2023.^44^ For this score, all genes in a cell type cluster (**Figure 1F**) were z-scored. The provided senescence gene list was split into genes that are up- and down-regulated in p16+ cells. The scale.data slot, which contains the z-scored counts, was extracted from the full dataset. The genes that were in the up- and down-regulated gene lists were identified and subset out of the extracted scale.data matrix. Two scores were calculated, one being the sum of the z-scored counts of the down-regulated genes multiplied by negative one and the other score being the sum of the z-scored counts of the up-regulated genes. The overall score is the sum of the down-regulated gene score and the up-regulated gene score scaled by the length of the gene set. With this method we tested six gene lists: Stimulation Independent, Replicative, Oncogene, Ion-Radiation Induced, Aged-Chondrocyte, and Two-way foreign body response (FBR). The gene lists were the result of differential expression analysis of bulk RNA-seq experiments comparing p16+ to p16– cultured fibroblasts (Stimulation Independent, Replicative, Oncogene, and Ion-Radiation Induced)^46^, uninjured aged chondrocytes (Aged-Chondrocytes) (B.O. Diekman, personal communication).^50–52^, and CD29+ cells from a FBR model (Two-way FBR)^44^.

The One-way Senescence Score was calculated using Single-Sample GSEA as per Saul et al., 2022.^47^ This method uses the raw counts of genes that are upregulated in p16+ cells. With this method we tested four gene lists: One-way FBR, SenMayo, GO: Senescence, and GO: SASP. The One-way FBR gene list contains only the upregulated genes found in the Two-way FBR gene list.^44^ The SenMayo gene list is a literature-curated list of genes commonly used to identify senescent cells.^47^ The GO: Senescence gene list is the GO: FRIDMAN_SENESCENCE_UP^48^ gene list and the GO: SASP gene list is the GO: REACTOME_SENESCENCE_ASSOCIATED_SECRETORY_PHENOTYPE_SASP^49^ gene list.

We evaluated the ability of the two methods and the ten gene lists to accurately identify senescent-like MuSCs and progenitors by calculating a receiver operator characteristic (ROC) curve. For each MuSCs and progenitor cell, we evaluated whether it expressed both *Cdkn2a* and *Cdkn1a*. For each method and gene list, the scores were ranked from highest to lowest and then grouped into 100 bins with approximately the same number of MuSCs and progenitors. We evaluated the specificity and the sensitivity in each bin where a true positive expresses both *Cdkn2a* and *Cdkn1a* and has a high senescent score. This analysis was done for all MuSCs and progenitors (**Figure 3M-N**), MuSCs and progenitors split by age (**Figure 3O**, Supplementary Figure 7A**-F**), and MuSCs and progenitors split by G1-Status (**Figure 5B**, Supplementary Figure 7G**-J**). The area under the curve (AUC) was calculated for each ROC curve (**Figures 3M**-O and 5B, Supplementary Figure 7K).

Given that the One-way Senescence Score with the FBR gene list performed the best (AUC = 0.86), we focused on that for further analyses. We next set a threshold of senescence based on the One-way FBR. We ranked the MuSCs and progenitors from highest to lowest One-way FBR Sen Score and then grouped the cells into 100 bins with approximately the same number of MuSCs and progenitors. Within each bin we calculated the fraction of MuSCs and progenitors that co-express *Cdkn2a* and *Cdkn1a*. The One-way FBR Sen Score where 50% of the MuSCs and progenitors with at least that score co-express *Cdkn2a* and *Cdkn1a* was set as the senescence threshold. For the One-way FBR Sen Score, MuSCs and progenitors with a score >= 2412.562 were called “senescent-like” while all other cells were called “not senescent-like” (**Figure 5A**, Supplementary Figure 11G**-H**).

#### Cell cycle scoring

To each cell in the final dataset, we assigned an S phase score, a G2M phase score, and a discrete phase classification (G1/S/G2M) using Seurat’s standard Cell-Cycling Scoring method.^64^ We have treated the S phase and G2M phase score as polar coordinates to help us visualize how cells are progressing through the cell cycle (**Figure 4D**). We have converted the polar coordinates to cartesian coordinates and normalized the theta values so that they range from 0 to 1 so that cells in G1 have the lowest theta values followed by cells in S and G2M (**Figure 4E**, Supplementary Figure 11A**-B**). This enables us to see how cells are progressing linearly through the cell cycle. Seurat’s standard cutoff between cells classified as G1 versus cells classified as S is at the normalized theta value of 0.25. Looking at the distribution of cells across the normalized theta values as well as the expression of cell cycle markers *Cdk1* and *Cdk4*, we decided to extend the G1 to S cutoff to 0.375 (**Figure 4E-G**, Supplementary Figure 11C**-D**). Cells with a normalized theta value >= 0.375 are considered Non-G1 (S/G2/M).

#### Myogenic cell subsets

From the final dataset, the cells with the cell type IDs ‘MuSCs and progenitors’ and ‘Myonuclei’ were subset out and the Seurat workflow was partially re-run (ScaleData, FindVariableFeatures, RunPCA, FindNeighbors, FindClusters, and RunUMAP). Dimensions accounting for 95% of the total variance were used to generate SNN graphs (FindNeighbors) and SNN clustering was performed (FindClusters).^64^ A clustering resolution of 0.7 was used resulting in 9 clusters. These 9 clusters were assigned general cell type IDs based on canonical myogenic markers. Of the 9 clusters, 4 were progenitor subtypes, 3 were myonuclei subtypes, and 2 were doublets (**Figure 4A**).

To more specifically ID the doublet clusters we looked at the co-expression of myogenic and non-myogenic markers (Supplementary Figure 8). Using GetAssayData, we extracted the log-normalized expression values of *Pax7*, *Pecam1*, *Acta1*, and *C1qa* in each cell in the 9 myogenic clusters. For each of the 9 clusters, we plotted cells by their expression values of *Pax7* and *Pecam1* and by their expression values of *Acta1* and *C1qa*. A density plot was plotted along the x- and y-axes using ggmarginal(type = “density”) (Supplementary Figure 8). The two clusters identified as doublets were excluded from the remaining myogenic subset analyses.

To identify the myonuclei clusters more specifically, we looked at the expression of myonuclei markers, markers of high metabolic activity, and the percent of unique transcripts derived from mitochondrial genes (Supplementary Figure 9). Using GetAssayData, we extracted the log-normalized expression values of *Myh1* and *Myh4* in each cell in the three myonuclei clusters. For every cell, as defined by the cell barcode, we determined whether the expression value equaled zero (no expression) or exceeded zero (expression) for both *Myh1* and *Myh4* independently. For each of the three myonuclei clusters, the fraction of cells that expressed *Myh1* and *Myh4*, only *Myh1*, only *Myh4*, and neither *Myh1* nor *Myh4* were calculate by dividing the number of cells that expressed *Myh1* and *Myh4*, only *Myh1*, only *Myh4*, and neither *Myh1* nor *Myh4* by the total number of cells within each myonuclei cluster (Supplementary Figure 9D**-G**).

Harmony embedding values from the dimensions accounting for 95% of the total variance were used for further dimensional reduction with PHATE, using phateR (v1.0.7) (**Figure 4B-C**).^60^ The PHATE embedding values were used by monocle3 (v1.0.0).^61–63^ The normal monocle3 workflow was used (cluster_cells, estimate_size_factors, learngraph, order_cells) where L1.sigma = 0.4 and the root cell was in the Progenitor 1 cluster. The pseudotime values for each cell as defined by monocle3 were transferred from the monocle3 CDS object to the myogenic cells only Seurat object by cell barcode. The pseudotime values were divided into 25 bins with approximately equal numbers of cells (1089-1090 cells per bin) (**Figure 4C**).

We assigned myogenic cell type IDs based on known myogenic marker expression in each pseudotime bin and dpi. We visualized this with dot plots where the size of the dot corresponds to the percent of cells in each pseudotime bin and dpi normalized by the dpi. The color of the dots corresponds to the average log-normalized expression of a select myogenic marker in each pseudotime bin and dpi (Supplementary Figure 10A**-H**). Cells in pseudotime bin 1 and dpi 0, 1, 2, 3.5, 5, and 7 were classified as ‘MuSCs 1’, cells in pseudotime bin 2 and dpi 0, 1, 2, 3.5, 5, and 7 were classified as ‘MuSCs 2’, cells in pseudotime bin 3 and dpi 0, 1, 2, 3.5, 5, and 7 were classified as ‘MuSCs 3’, cells in pseudotime bin 4 and dpi 0, 1, 2, 3.5, 5, and 7 were classified as ‘MuSCs 4’, cells in pseudotime bin 5 and dpi 0, 1, 2, 3.5, 5, and 7 were classified as ‘MuSCs 5’, and cells in pseudotime bin 6 and dpi 0, 1, 2, 3.5, 5, and 7 and cells in pseudotime bin 7 and dpi 5 and 7 were classified as ‘MuSCs 6’ based on expression of *Pax7* and *Myf5* (Supplementary Figure 10A**-B**,I). Cells in pseudotime bins 7-13 and dpi 0 were classified as ‘Non-Cycling MPCs’ based on the expression of *Pax7*, *Myf5*, and *Myod1* and the lack of expression of cycling markers *Cdk1* and *Cdk4* (Supplementary Figure 10A**-C**,G-I). Cells in pseudotime bin 7 and dpi 1, 2, and 3.5 and cells in pseudotime bins 8-11 and dpi 1, 2, 3.5, 5, and 7 were classified as ‘Cycling MPCs’ based on the expression of *Myod1* and cycling markers *Cdk1* and *Cdk4* (Supplementary Figure 10C**,G**-I). Cells in pseudotime bin 12 and dpi 0, 1, 2, 3.5, 5, and 7 and cells in pseudotime bin 13 and dpi 1 and 2 were classified as ‘Committing MPCs’ based on the low expression of *Myod1*, *Myog*, *Mymk* and the still high expression of cycling markers *Cdk1* and *Cdk4* (Supplementary Figure 10C**-E**,G-I). Cells in pseudotime bin 13 and dpi 3.5, 5, and 7 were classified as ‘Fusing Myocytes’ based on the high expression of *Myog* and Mymk and low expression of cycling markers *Cdk1* and *Cdk4* (Supplementary Figure 10D**-E**,G-I). Cells in pseudotime bins 14-25 and dpi 0, 1, 2, 3.5, 5, and 7 were classified as ‘Myonuclei’ based on the expression of *Acta1* (Supplementary Figure 10F**,I**).

We focused on dpi 3.5 and compared the cells in MuSCs 1, MuSCs 2-6, and all MPCs (this includes cells classified as Cycling MPCs and cells classified as Committing MPCs). For each sample, we calculated the fraction of cells that had log-normalized *Cdkn2a* and *Cdkn1a* counts greater than 0 (**Figure 5E-G**). Within these same groupings we also calculated the fraction of cells that had a One-Way FBR score greater than 2412.562 (**Figure 6H-J**). We conducted unpaired, two-sided Student’s t-tests to evaluate whether there was a significant difference between ages.

#### Differential expression analysis and pre-ranked Gene Set Enrichment Analysis

For select comparisons we used Seurat’s FindAllMarkers() function to identify genes that were differentially expressed between groups. In the myogenic subset with all ages at day 3.5, we did this analysis between the cells in ‘MuSCs 1’ and the cells in ‘MuSCs 2-6’ (**Figure 4L**). In the MuSCs and progenitors with all ages at day 3.5, we did this analysis between cells that co-expressed *Cdkn2a* and *Cdkn1a* (we refer to these cells as ‘Double positive’) and all other cells (we refer to these cells as ‘other’). Genes that had an FDR-corrected p-value <=0.05 were ranked by average log2 fold change and used in a Gene Set Enrichment Analysis (GSEA, v4.1.0). The gene set databases used included h.all.v2023.1.Hs.symbols.gmt, c2.all.v2023.1.Hs.symbols.gmt, 5.all.v2023.1.Hs.symbols.gmt, and c8.all.v2023.1.Hs.symbols.gmt. Significant GO Terms (FDR q-value <= 0.25) were ranked by the normalized enrichment score (enrichment scores normalized by the size of the gene set) (**Figure 3H**).

#### Key resource availability

A complete list of metadata and GEO accessions for the scRNA-seq data can be found in Extended Data File 1. Previously published scRNA-seq data are deposited in GEO under accessions GSE143437, GSE159500, and GSE162172. Newly collected scRNA-seq data from young (4.7 mo; days 1 and 3.5) and geriatric (26 mo) mice are deposited in GEO under accession GSE232106. Fully processed Seurat objects for the final dataset (**Figure 1**) and the myogenic subset (**Figures 4** and **5**) will be available for download on Dryad upon publication. Gene lists used in the Senescence Scoring analysis are compiled in the Extended Data File 2. All newly developed code (Senescence Scoring, Cell Cycle Scoring, Pseudotime Binning) will be available on Github upon publication.

## AUTHOR CONTRIBUTIONS

L.D.W., V.I.M., B.D.R., J.H.E., and B.D.C. designed the study. L.D.W., J.L.O., E.H.H.F., and V.I.M. carried out the experiments. L.D.W. performed computational analyses. L.D.W. and J.L.O. analyzed experimental data and prepared figures. B.D.R., J.H.E., and B.D.C. supervised the data analyses. L.D.W. and B.D.C. wrote the manuscript. All authors provided feedback and comments.

## ACKNOWLEDGEMENTS

We thank Peter Schweitzer and colleagues in the Genomics Facility (Research Resource Identifier RRID:SCR_021727) of the Cornell Biotechnology Resource Center in the Cornell Institute for Biotechnology for their help in performing single-cell RNA-sequencing experiments. We thank Lydia Tesfa and colleagues in the Flow Cytometry Facility (RRID:SCR_021740) of the Cornell Biotechnology Resource Center for their help in performing flow cytometry experiments. We thank the Cornell Center for Animal Resources and Education for assisting in animal housing and care. We thank Andrea De Micheli and Ern Hwei Hannah Fong for helping with mouse procedures and in generating some single-cell RNA sequencing data. We thank Christopher Cherry in the Elisseeff research group and other members of the JHU-Mayo-NIH SenNet team for advice on senescence scoring methods. We thank Brian Diekman and his research group for generating and providing the chondrocyte gene signature. We thank David McKellar for providing code via his Github repository and helpful advice. This work was supported by the US National Institutes of Health (NIH) grants R01AG058630 (to B.D.C.), U54AG07977 (to B.D.C. and J.H.E.), T32HD057854 (to L.D.W.), F30OD032097 (to V.I.M.), R01AI105265 (to B.D.R.), DP1AR076959 (to J.H.E.). The content is solely the responsibility of the authors and does not necessarily represent the official views of the NIH. We also acknowledge funding support from the Bloomberg∼Kimmel Institute (to J.H.E.) and Morton Goldberg Professorship (to J.H.E.).

---

**Supplemental Figure 1:**
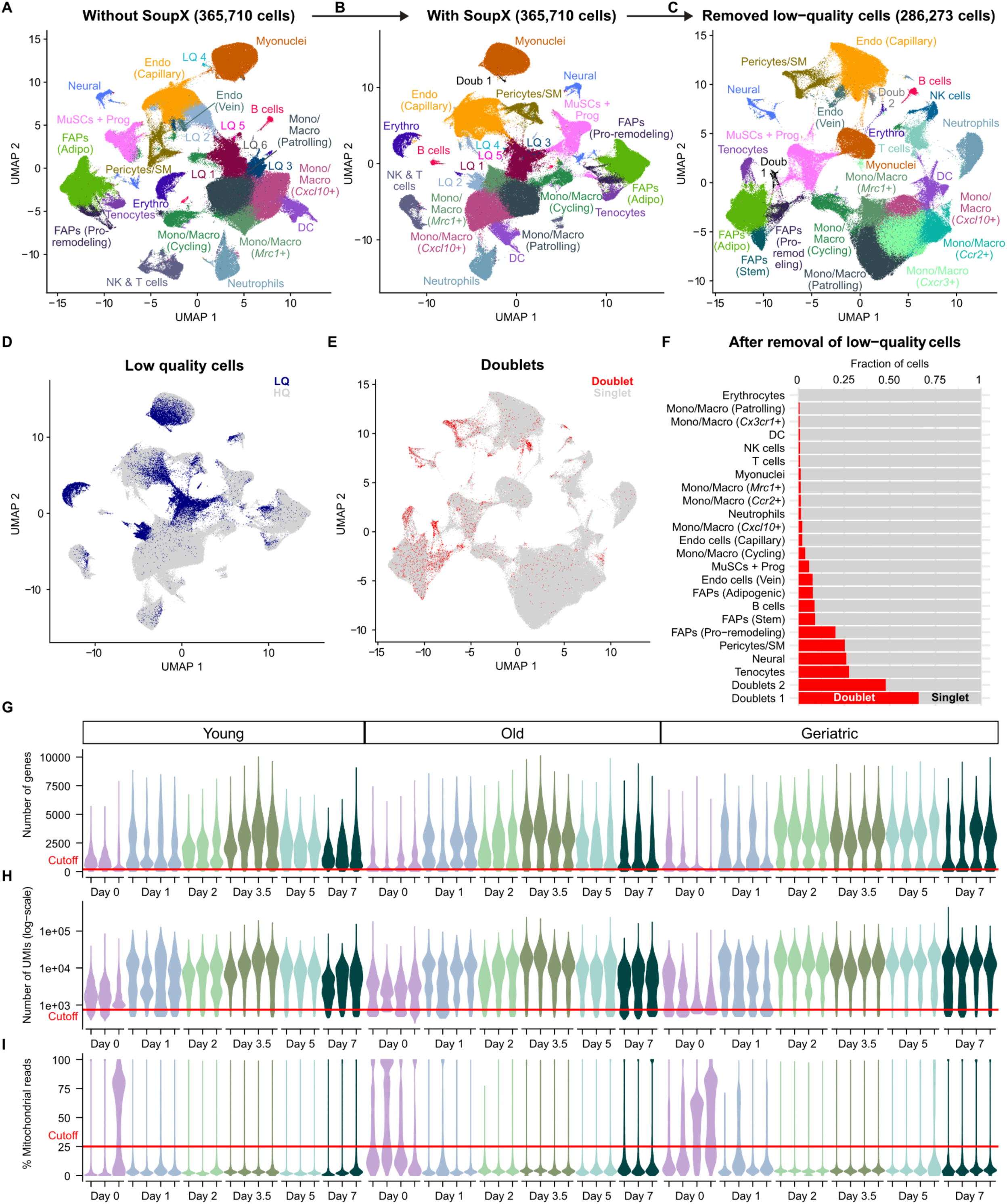
Evaluating sample quality. (**A-C**) Quality control workflow and resulting UMAPs with cells colored by manually assigned cell type IDs based on the expression of hallmark skeletal muscle genes. Prior to any quality-control there were 365,710 cells and 24 cell types were identified, including 6 low-quality (LQ) clusters (**A**). After ambient RNA removal with SoupX, 23 cell types were identified, including 5 LQ clusters (**B**). After ambient RNA removal with SoupX and removal of LQ cells based on the number of genes and UMIs and the percent of mitochondrial reads, there were 286,273 cells and 24 cell types were identified, including two doublet clusters (**C**). (**D**) This is the same UMAP as in (**B**), but the cells are colored by quality status. Cells that had <200 genes, <750 UMIs, and >25% mitochondrial reads are considered LQ. All other cells are considered high-quality (HQ). (**E**) This is the same UMAP as in (**C**), but the cells are colored by doublet status as determined by DoubletFinder using an estimated doublet rate of 5%. (**F**) For every cell type cluster in (**C**), the fraction of singlets and doublets was calculated. (**G-I**) Violin plots of the number of genes (**G**), the number of UMIs (**H**), and the percent of mitochondrial reads (**I**) in each sample.

**Supplemental Figure 2:**
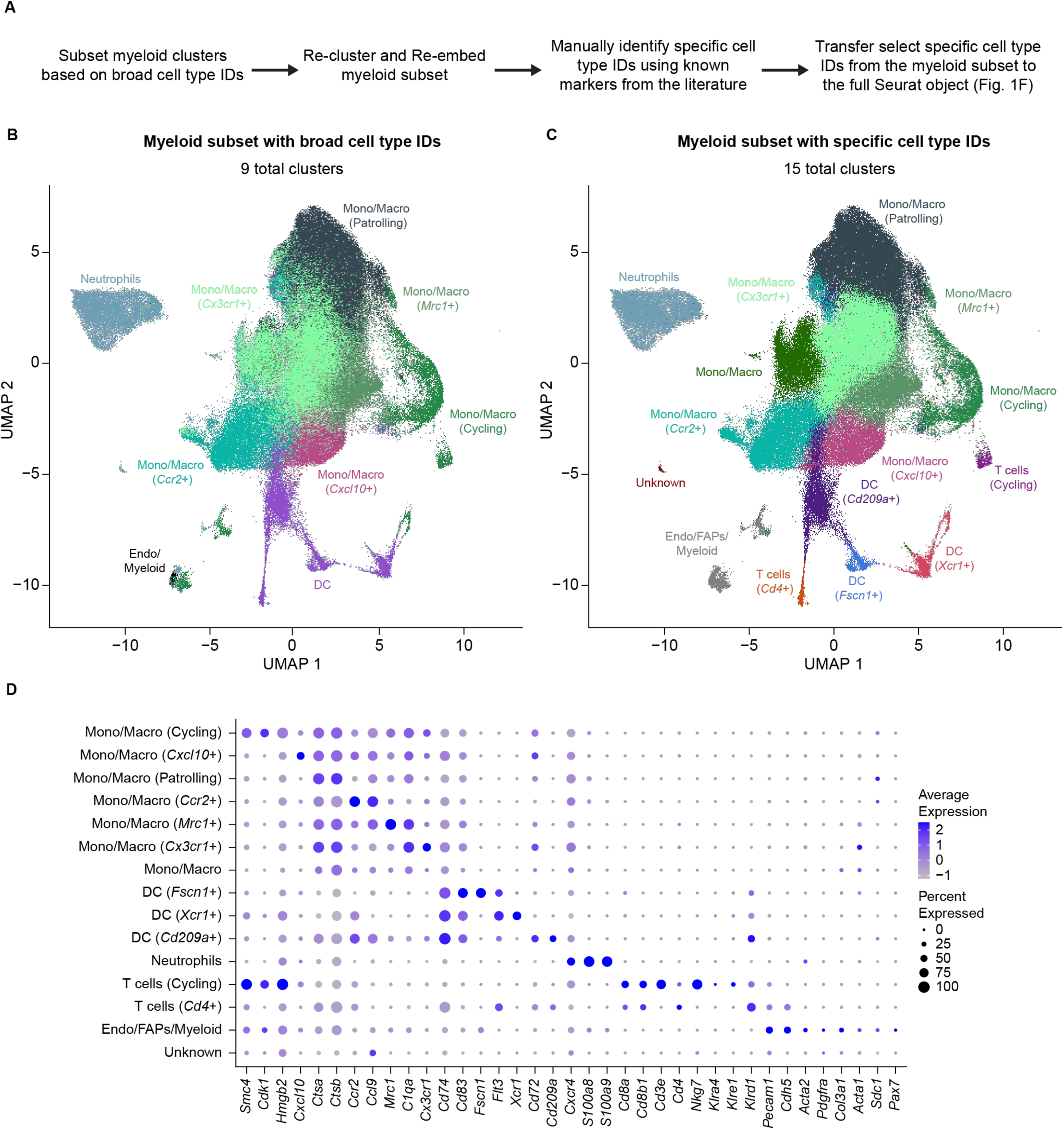
Identifying additional immune cell types when focusing on just the myeloid cells. (**A**) Workflow to clarify the myeloid cell types. The 9 myeloid clusters based on broad cell type IDs were subset from the final dataset, re-clustered, and re-embedded. More specific cell type IDs were manually assigned using genes known to mark myeloid cell types. Some of these more specific cell type IDs from the myeloid subset were transferred back to the final dataset (Figure 1F) based on the cell barcodes. (**B-C**) UMAPs of the myeloid subset after re-embedding, the cells are colored by the broad cell type IDs originally identified in the final dataset (**B**). After re-clustering, re-embedding, and re-annotating based on known immune cell markers, the cells are colored by the more specific cell type IDs (**C**). If the same cell type is in (**B**) and (**C**), it has the same color designation in both UMAPs. (**D**) Dot plot showing the expression frequency and magnitude of genes used to manually assign the more specific cell type IDs as shown in (**C**).

**Supplemental Figure 3:**
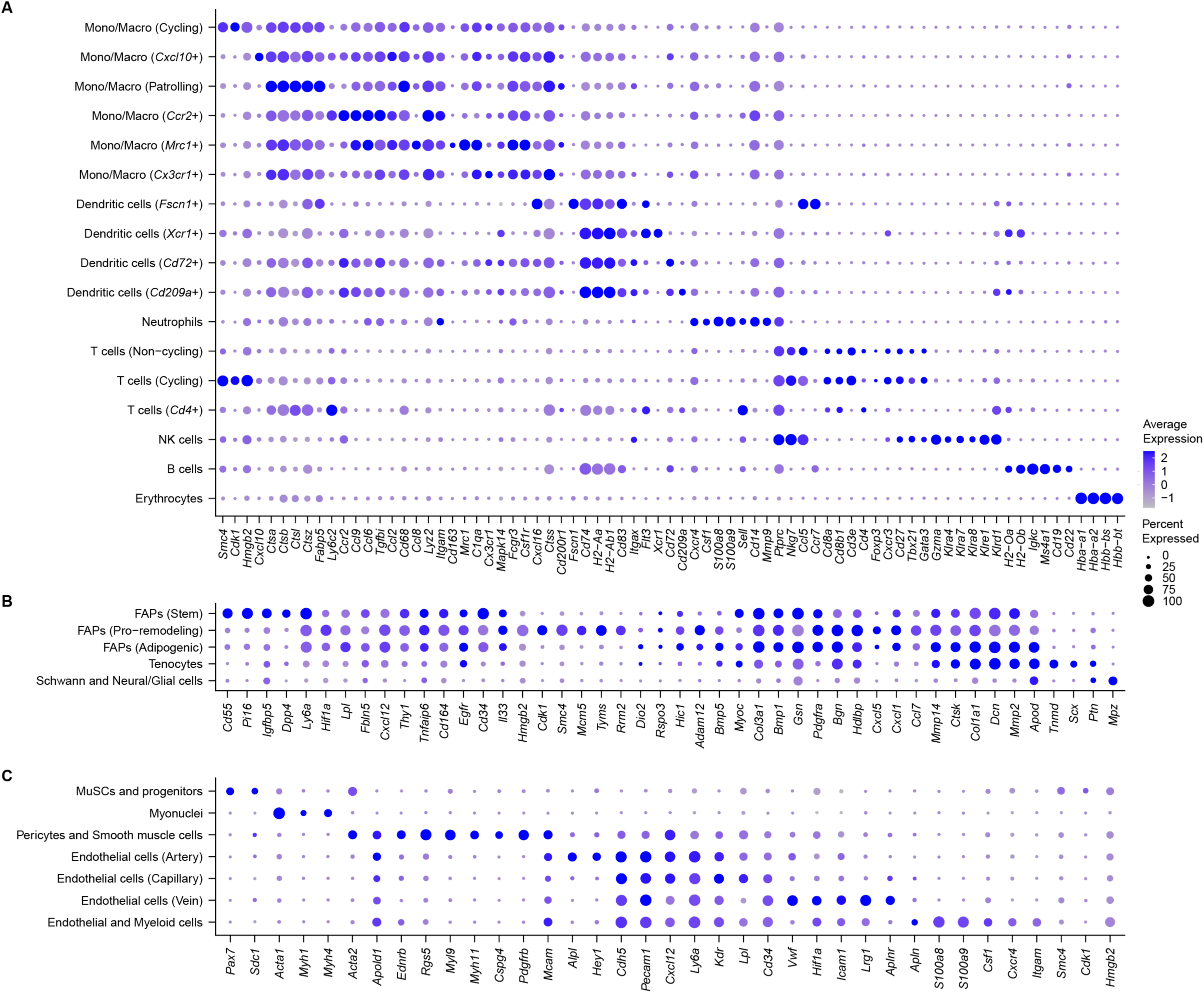
Identifying cell types in skeletal muscle during homeostasis and regeneration. (**A-C**) Dot plots showing the expression frequency and magnitude of genes used to manually assign cell type IDs, including the more specific immune cell type IDs. We identified 17 immune cell clusters (**A**), 5 FAPs, Tenocytes, and Neural cell clusters (**B**), and 7 myogenic, pericyte/skeletal muscle, and endothelial cell clusters (**C**).

**Supplemental Figure 4:**
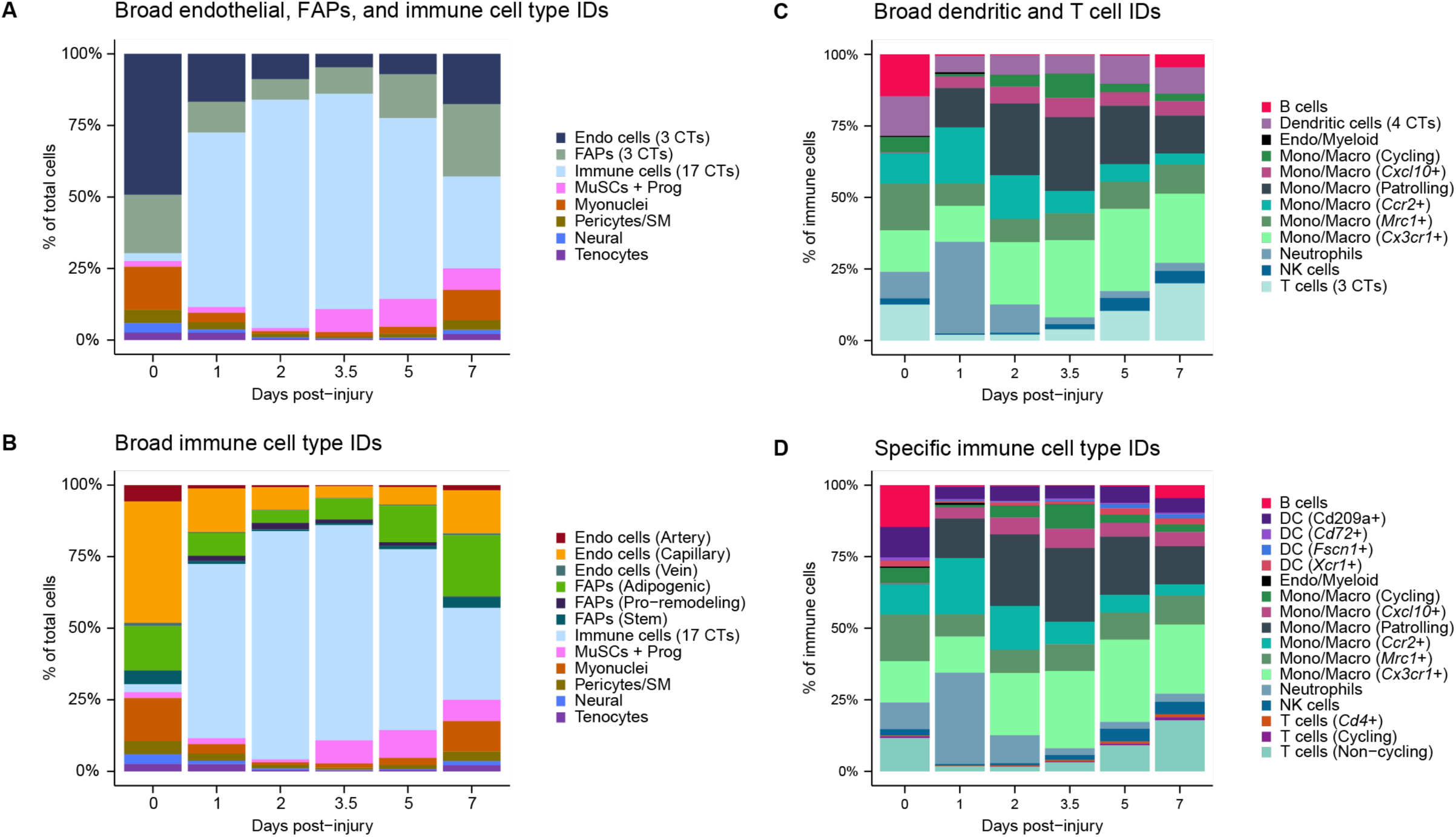
Determining the cell type composition in each day post injury. (**A-B**) Stacked bar plots of the percent of each cell type found at a given day post injury (dpi). The three endothelial populations were combined into “Endo (3 CTs)”, the three FAPs populations were combined into “FAPs (3 CTs)”, and the 17 immune populations were combined into “Immune cells (17 CTs)” (**A**). Only the 17 immune populations are combined into “Immune cells (17 CTs)” (this plot is the same as in Figure 2K) (**B**). (**C-D**) Stacked bar plots of the percent of each immune cell type out of all immune cells at a given dpi. The four dendritic cell populations were combined into “Dendritic cells (4 CTs)” and the three T cell populations were combined into “T cells (3 CTs)” (C). All individual immune cell type IDs (**D**).

**Supplemental Figure 5:**
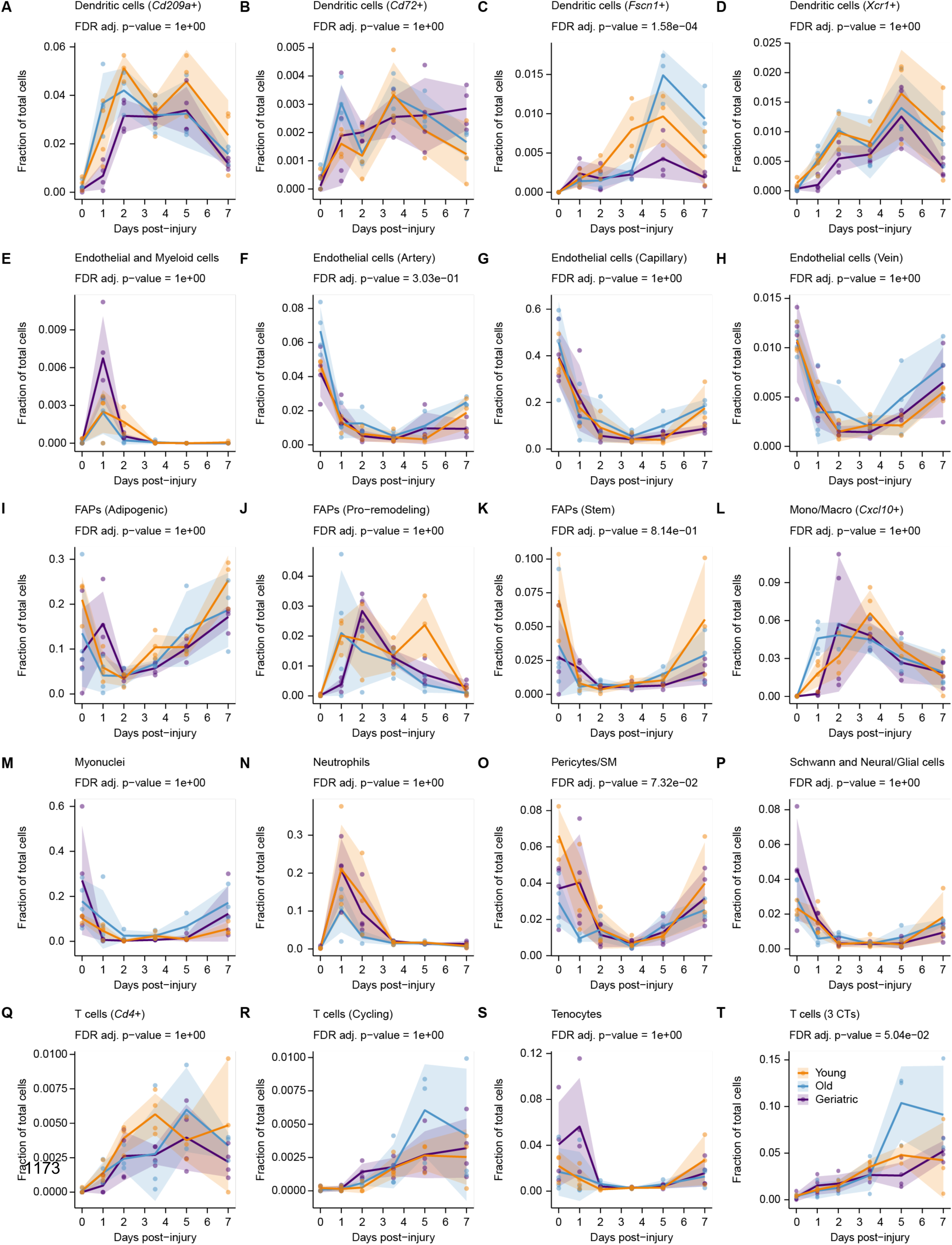
Cell type dynamics (those that are not in main. Figure 2). (**A-T**) Line plots for the remaining cell types not shown in Figure 2 showing cell type relative abundance as a fraction of total cells from 0-7 days post injury (dpi). For each sample, the number of cells of the reported type was divided by the total number of cells (excluding erythrocytes). Points are each sample (n = 3-4). Ribbon is the standard deviation. Statistical significance of age-specific cell type dynamics was evaluated using non-linear modeling and FDR-corrected p-values are reported (see **Supplementary** Figure 6).

**Supplemental Figure 6:**
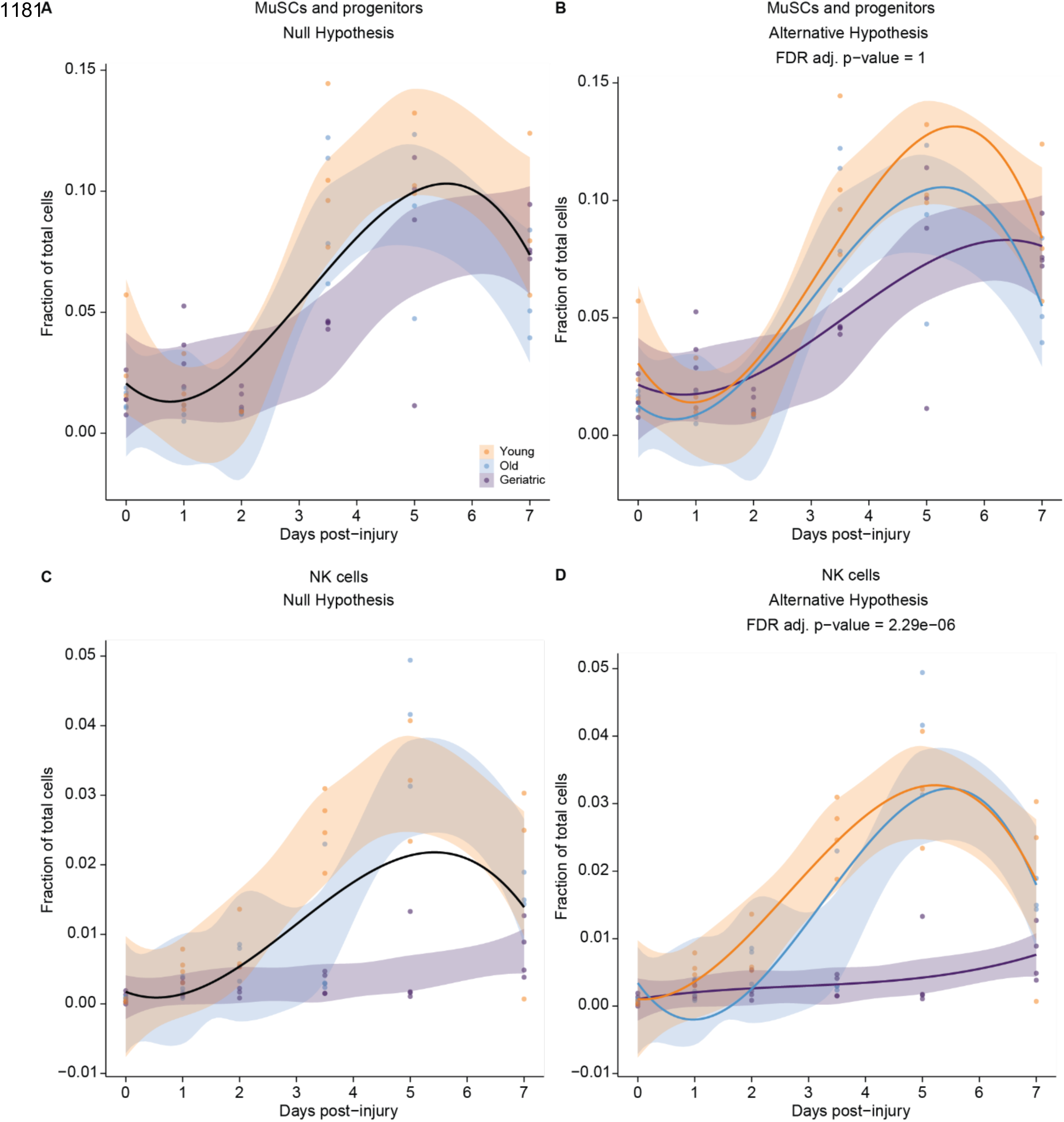
Cell type dynamics non-linear modeling. (**A-B**) Plots of the fraction of MuSCs and progenitors from 0-7 days post injury (dpi) by age. For each sample, the number of cells was divided by the total number of cells (excluding erythrocytes). The points are the fraction for each sample (n = 3-4) and the ribbon is the confidence interval. The black line is the non-linear model independent of age (**A**) and the three colored lines are the non-linear model dependent on age (**B**). Whether there was a significant difference in the non-linear models independent of and dependent on age was determined using a likelihood ratio test (ANOVA) and the p-values were corrected with FDR. (**C-D**) Same as in (**A-B**) but with NK cells.

**Supplemental Figure 7:**
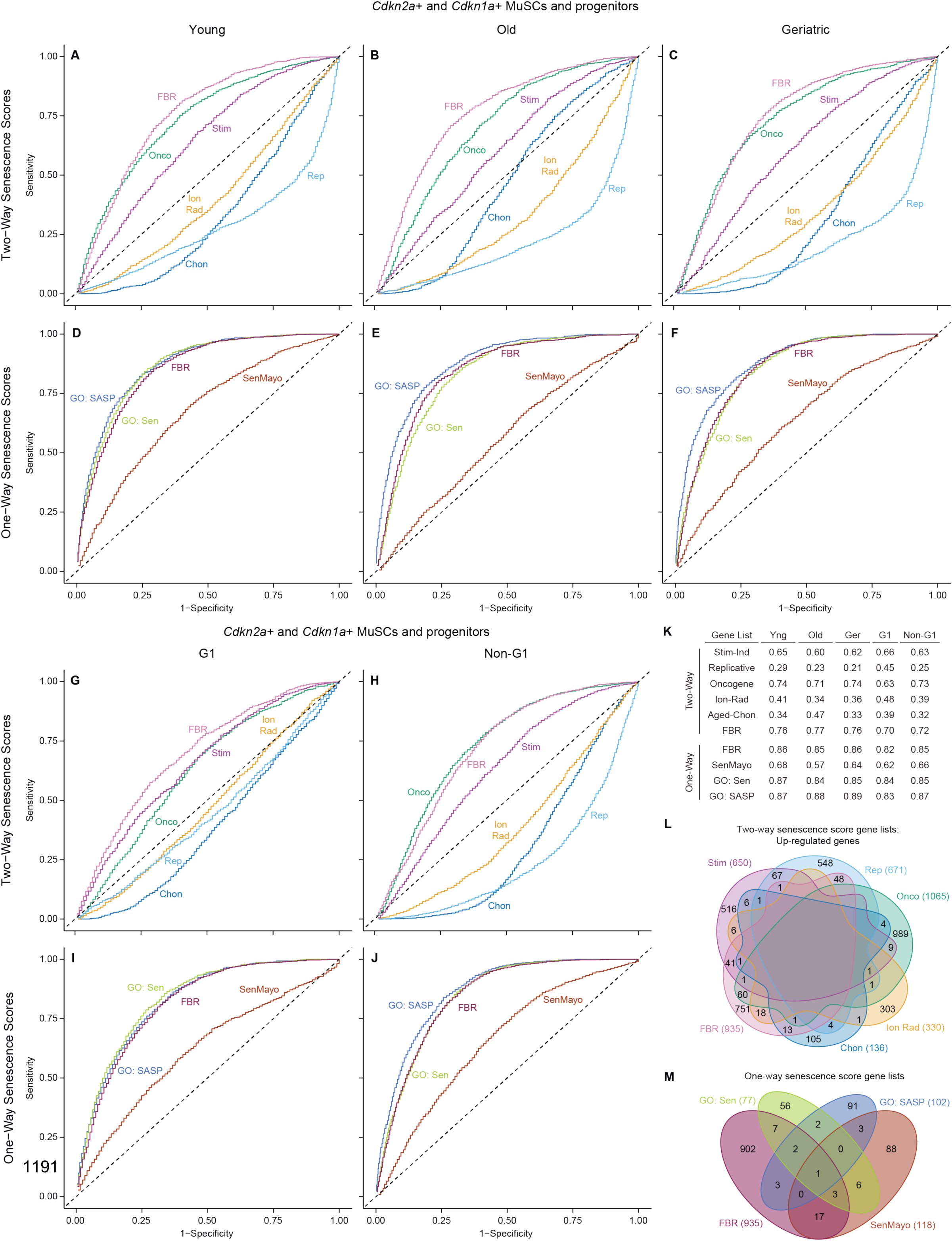
Evaluation of accuracy of senescence scoring methods. (**A-F**) Receiver Operator Characteristic (ROC) curves based on the co-expression of *Cdkn2a* and *Cdkn1a* for the six Two-way senescence scores (**A-C**) and the four One-way senescence scores (**D-F**) split into young (**A, D**), old (**B, E**), and geriatric (**C, F**) MuSCs and progenitors. (**G-J**) ROC curves based on the co-expression of *Cdkn2a* and *Cdkn1a* for the six Two-way senescence scores (**G-H**) and the four One-way senescence scores (**I-J**) split into G1 (**G, I**) and non-G1 (S, G2, and M) (**H, J**) MuSCs and progenitors. (**K**) Table of the area under the curve (AUC) for each ROC curve. (**L-M**) Venn diagrams of the unique and shared up-regulated genes found in the Two-way senescence scores (**L**) and the One-way senescence scores (**M**).

**Supplemental Figure 8:**
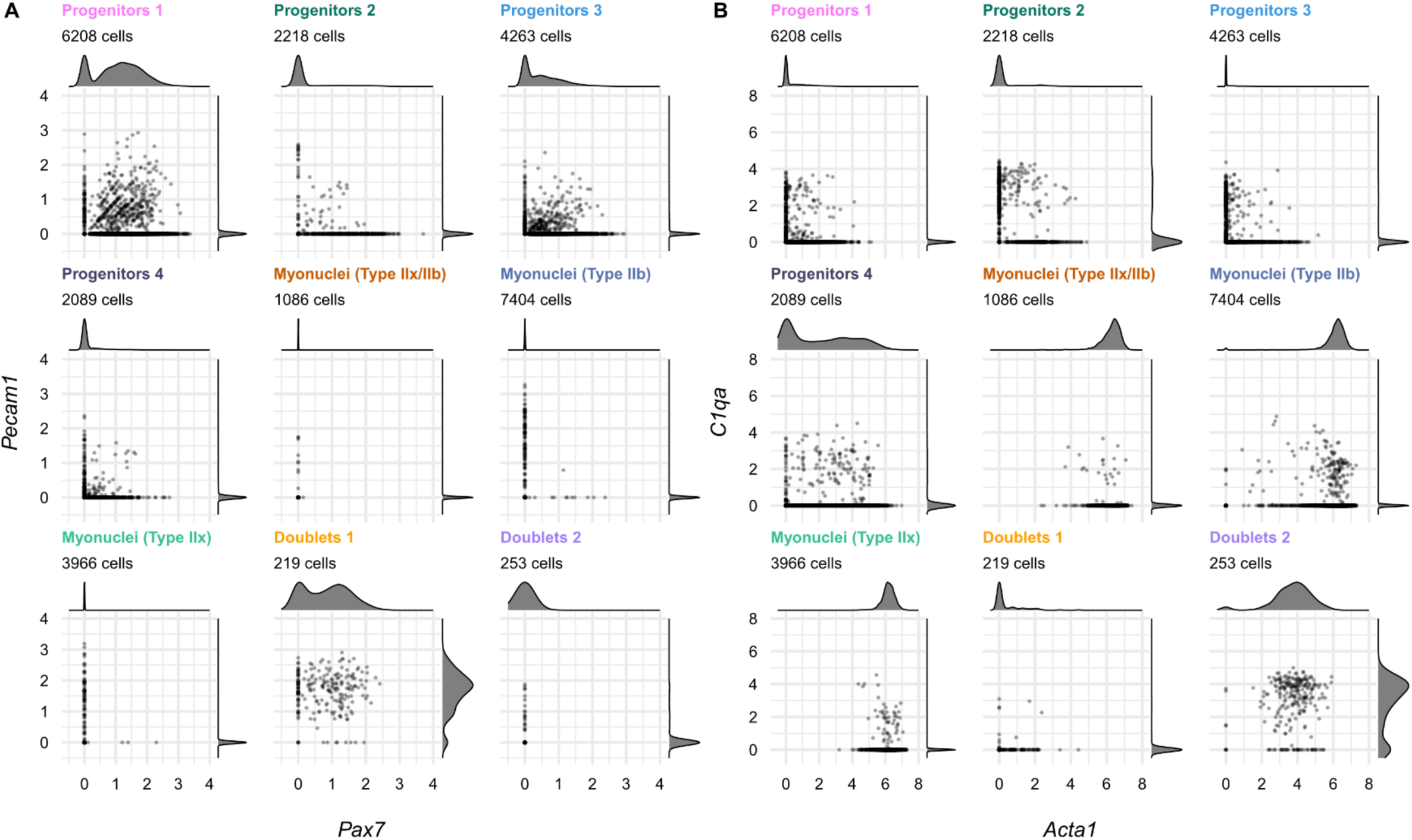
Identified two doublet clusters in the myogenic subset that were not identified by DoubletFinder. (**A-B**) Scatter plots of the log-normalized expression levels of *Pax7* and *Pecam1* (**A**) and *Acta1* and *C1qa* (**B**) by broad myogenic IDs as defined in Figure 4A. A density curve is plotted on each axis.

**Supplemental Figure 9:**
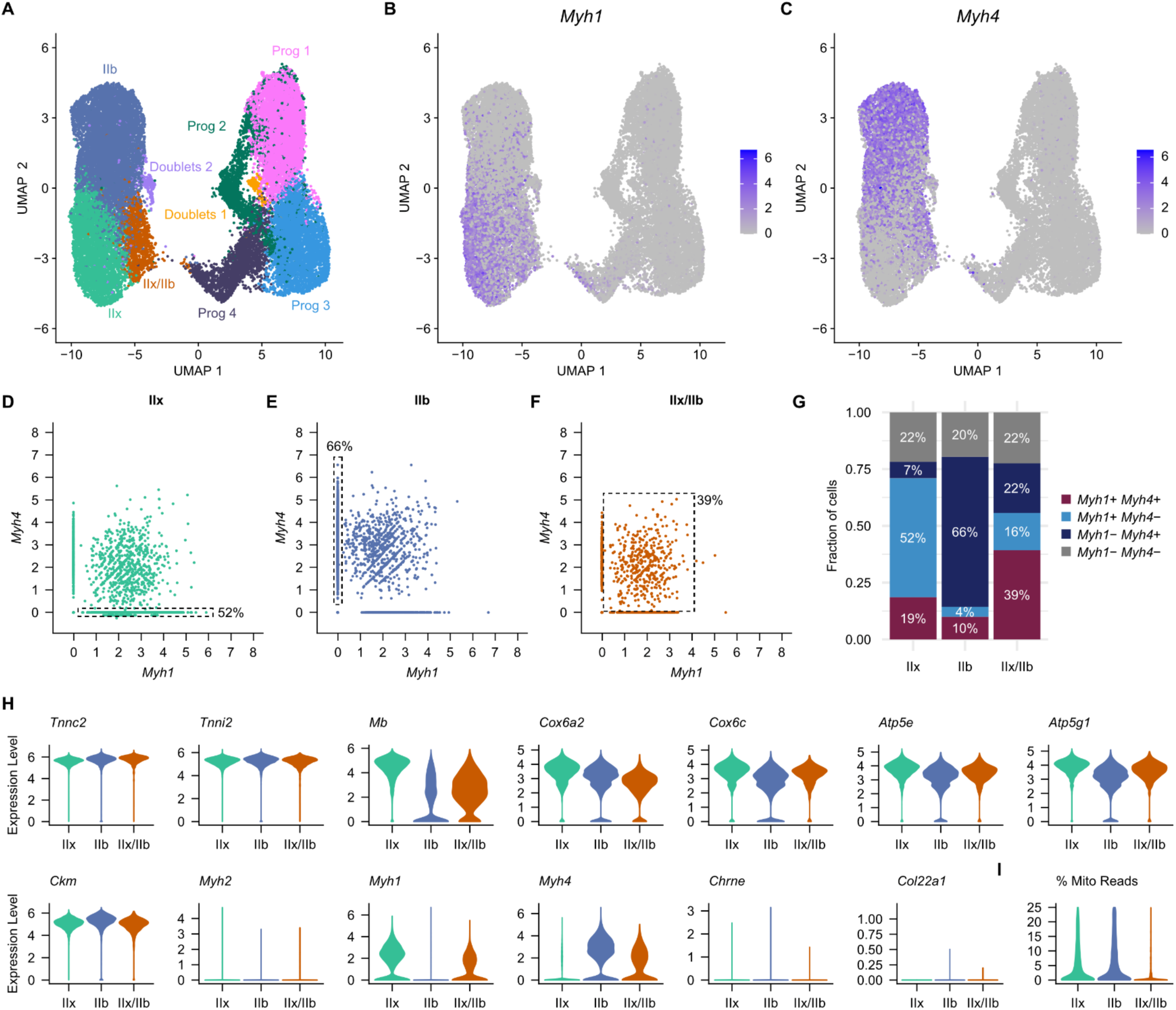
Identifying a transitional myonuclei state. (**A-C**) Same UMAP of the myogenic subset as in Figure 4A colored by broad myogenic IDs (**A**) and colored by the log-normalized expression of *Myh1* (**B**) and *Myh4* (**C**). (**D-F**) Scatter plots of the log-normalized expression levels of *Myh1* and *Myh4* in the myonuclei clusters Type IIx (**D**), Type IIb (**E**), and Type IIx/IIb (**F**). The dotted-line boxes highlight the cells that express the marker(s) that should be expressed in each myonuclei cluster. (**G**) Stacked bar plot of the fraction of cells within each myonuclei cluster that express a combination of *Myh1* and *Myh4*. (**H**) Violin plots of the log-normalized expression levels of markers of high metabolic rate (*Tnnc2*, *Tnni2*, *Mb*, *Cox6a2*, *Cox6c*, *Atp5e*, *Atp5g1*) and myonuclei (*Ckm*, *Myh2*, *Myh1*, *Myh4*, *Chrne*, *Col22a1*) in each myonuclei cluster. (**I**) Violin plot of the percent of mitochondrial reads in each myonuclei cluster.

**Supplemental Figure 10:**
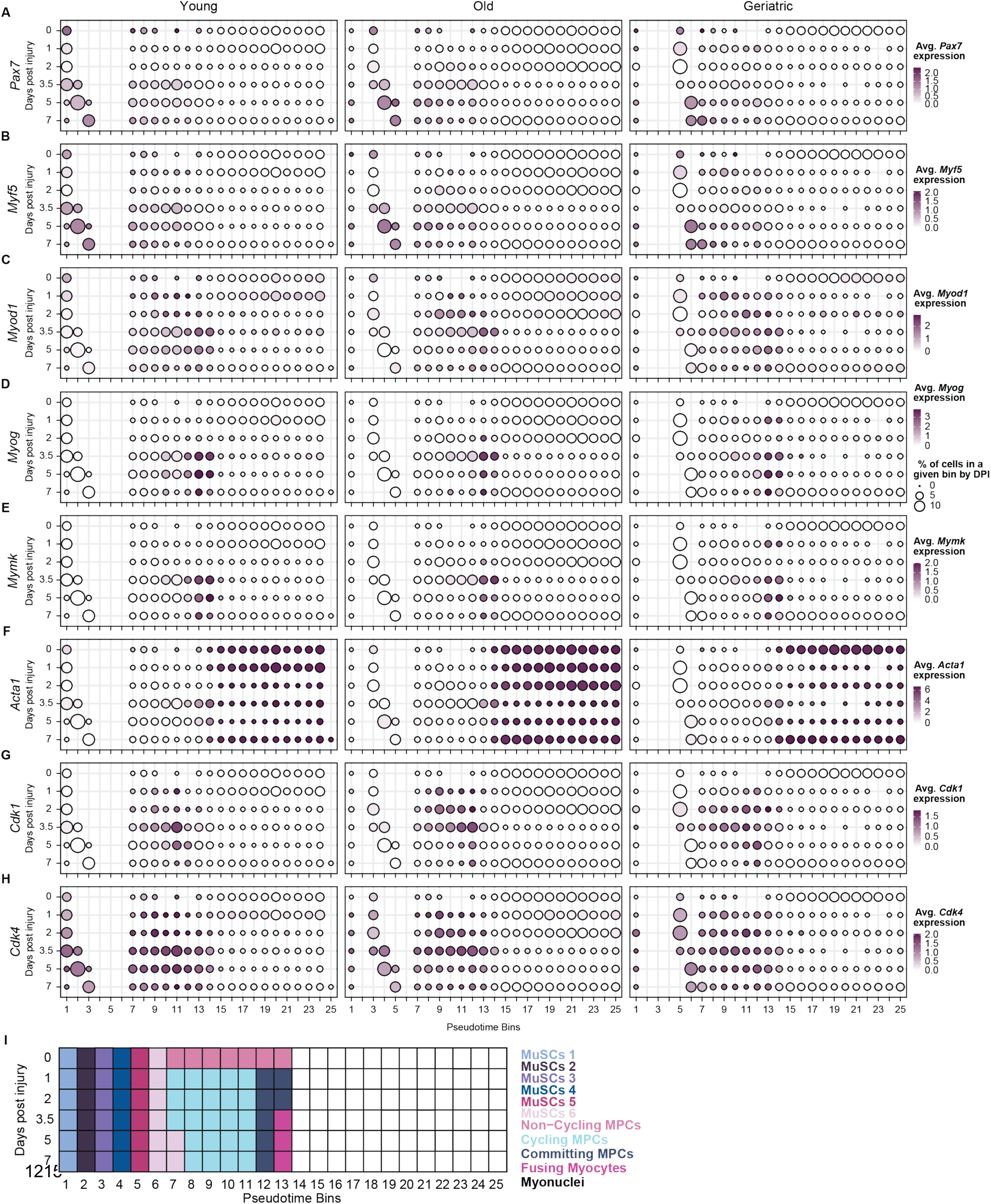
Myogenic subset pseudotime analysis. (**A-H**) Dot Plots of the average log-normalized expression of *Pax7* (**A**), *Myf5* (**B**), *Myod1* (**C**), *Myog* (**D**), *Mymk* (**E**), *Acta1* (**F**), *Cdk1* (**G**), and *Cdk4* (**H**) in each day post injury (dpi) and pseudotime bin. The size of the circle is the percent of cells in each pseudotime bin for each age and dpi combination. (**I**) Schematic of newly assigned myogenic IDs (referred to as ‘pseudotime-based myogenic cell state bin’) based on the expression of known myogenic markers in each dpi and pseudotime bin.

**Supplemental Figure 11:**
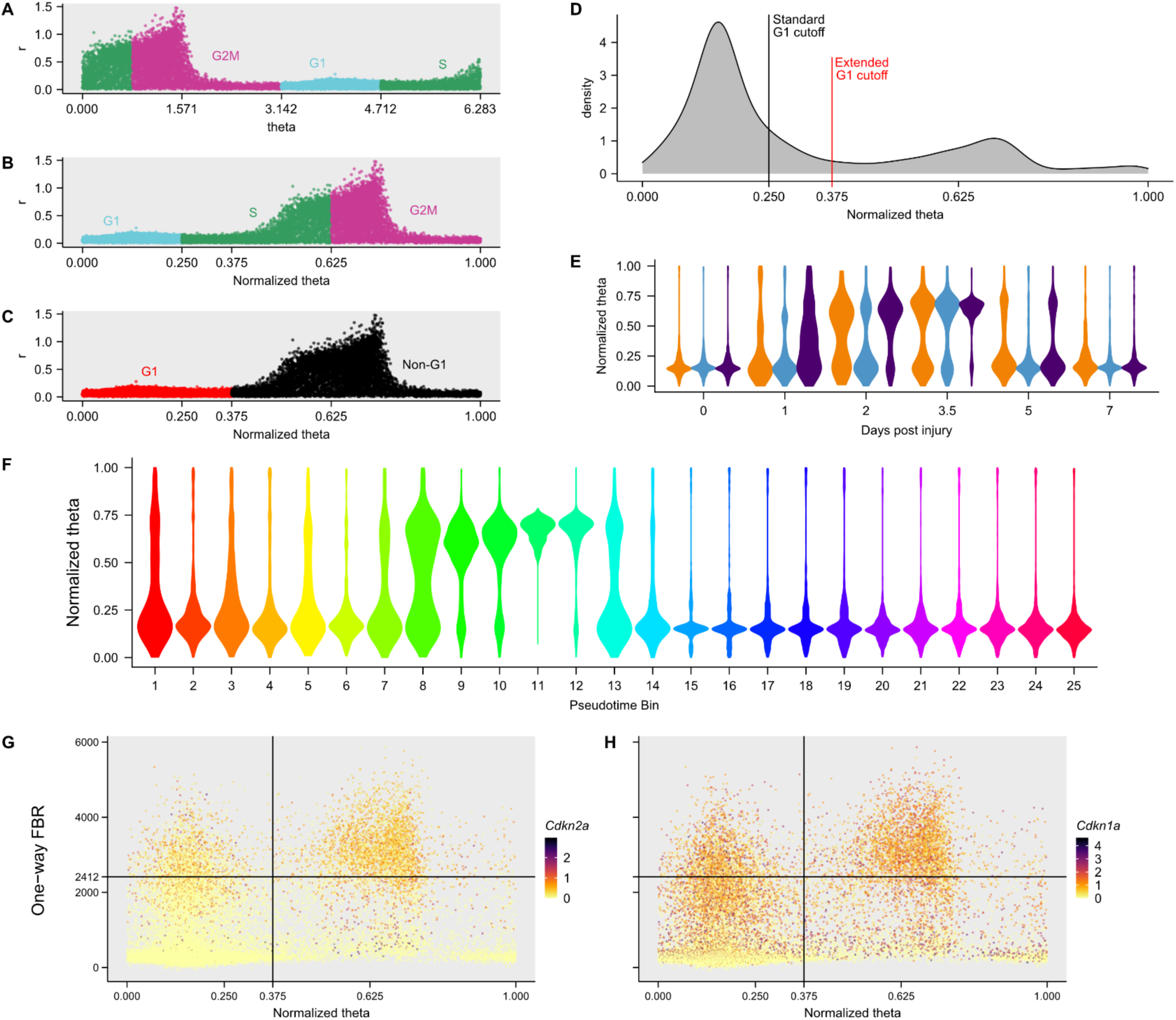
Cell cycle workflow. (**A-C**) The polar coordinates in Figure 4D were converted to cartesian coordinates (**A**) and rescaled to fit in a range from 0 to 1 (this plot is the same as in Figure 4E) (**B**). The cells are colored by the predicted cell cycle phase based on the two scores (**A-B**) and G1 status (**C**). (**D**) Distribution of MuSCs and progenitors across the normalized theta values. The black line represents the standard Seurat G1 cutoff (normalized theta = 0.25), and the red line represents our extended G1 cutoff (normalized theta = 0.375). (**E**) Violin plot of the distribution of the normalized theta values in MuSCs and progenitors split by age and days post injury (dpi). (**F**) Violin plot of the distribution of the normalized theta values in MuSCs and progenitors split by pseudotime bin. (**G-H**) Scatter plot of the normalized theta values and the One-way FBR senescence score in MuSCs and progenitors. The cells are colored by log-normalized *Cdkn2a* expression (**G**) and by *Cdkn1a* expression (**H**). The vertical line is the extended G1 cutoff, and the horizontal line is where 50% of the cells above this line co-express *Cdkn2a* and *Cdkn1a*.

**Supplementary Table 1:**
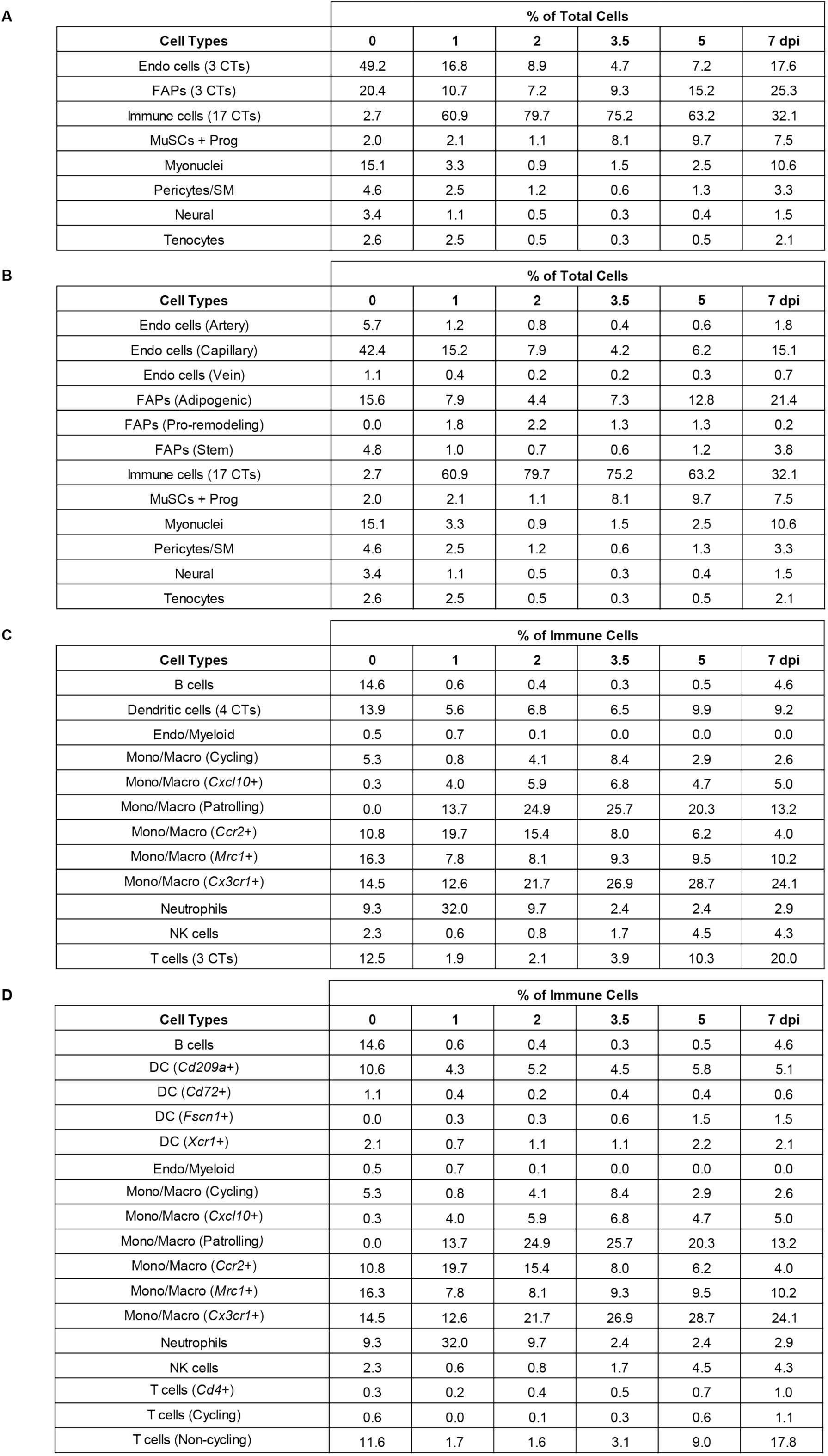
**Determining the cell type composition in each day post injury.** (**A-B**) Percent of each cell type found at a given day post injury (dpi). The three endothelial populations were combined into “Endo (3 CTs)”, the three FAPs populations were combined into “FAPs (3 CTs)”, and the 17 immune populations were combined into “Immune cells (17 CTs)” (**A**). Only the 17 immune populations are combined into “Immune cells (17 CTs)” (**B**). (**C-D**) Percent of each immune cell type out of all immune cells at a given dpi. The four dendritic cell populations were combined into “Dendritic cells (4 CTs)” and the three T cell populations were combined into “T cells (3 CTs)” (**C**). All individual immune cell type IDs (**D**).

| Cell Types | % of Total Cells |  |  |  |  |  |
| --- | --- | --- | --- | --- | --- | --- |
|  | 0 | 1 | 2 | 3.5 | 5 | 7 dpi |
| Endo cells (3 CTs) | 49.2 | 16.8 | 8.9 | 4.7 | 7.2 | 17.6 |
| FAPs (3 CTs) | 20.4 | 10.7 | 7.2 | 9.3 | 15.2 | 25.3 |
| Immune cells (17 CTs) | 2.7 | 60.9 | 79.7 | 75.2 | 63.2 | 32.1 |
| MuSCs + Prog | 2.0 | 2.1 | 1.1 | 8.1 | 9.7 | 7.5 |
| Myonuclei | 15.1 | 3.3 | 0.9 | 1.5 | 2.5 | 10.6 |
| Pericytes/SM | 4.6 | 2.5 | 1.2 | 0.6 | 1.3 | 3.3 |
| Neural | 3.4 | 1.1 | 0.5 | 0.3 | 0.4 | 1.5 |
| Tenocytes | 2.6 | 2.5 | 0.5 | 0.3 | 0.5 | 2.1 |

B
| Cell Types | % of Total Cells |  |  |  |  |  |
| --- | --- | --- | --- | --- | --- | --- |
|  | 0 | 1 | 2 | 3.5 | 5 | 7 dpi |
| Endo cells (Artery) | 5.7 | 1.2 | 0.8 | 0.4 | 0.6 | 1.8 |
| Endo cells (Capillary) | 42.4 | 15.2 | 7.9 | 4.2 | 6.2 | 15.1 |
| Endo cells (Vein) | 1.1 | 0.4 | 0.2 | 0.2 | 0.3 | 0.7 |
| FAPs (Adipogenic) | 15.6 | 7.9 | 4.4 | 7.3 | 12.8 | 21.4 |
| FAPs (Pro-remodeling) | 0.0 | 1.8 | 2.2 | 1.3 | 1.3 | 0.2 |
| FAPs (Stem) | 4.8 | 1.0 | 0.7 | 0.6 | 1.2 | 3.8 |
| Immune cells (17 CTs) | 2.7 | 60.9 | 79.7 | 75.2 | 63.2 | 32.1 |
| MuSCs + Prog | 2.0 | 2.1 | 1.1 | 8.1 | 9.7 | 7.5 |
| Myonuclei | 15.1 | 3.3 | 0.9 | 1.5 | 2.5 | 10.6 |
| Pericytes/SM | 4.6 | 2.5 | 1.2 | 0.6 | 1.3 | 3.3 |
| Neural | 3.4 | 1.1 | 0.5 | 0.3 | 0.4 | 1.5 |
| Tenocytes | 2.6 | 2.5 | 0.5 | 0.3 | 0.5 | 2.1 |

C
| Cell Types | % of Immune Cells |  |  |  |  |  |
| --- | --- | --- | --- | --- | --- | --- |
|  | 0 | 1 | 2 | 3.5 | 5 | 7 dpi |
| B cells | 14.6 | 0.6 | 0.4 | 0.3 | 0.5 | 4.6 |
| Dendritic cells (4 CTs) | 13.9 | 5.6 | 6.8 | 6.5 | 9.9 | 9.2 |
| Endo/Myeloid | 0.5 | 0.7 | 0.1 | 0.0 | 0.0 | 0.0 |
| Mono/Macro (Cycling) | 5.3 | 0.8 | 4.1 | 8.4 | 2.9 | 2.6 |
| Mono/Macro (*Cxcl10*+) | 0.3 | 4.0 | 5.9 | 6.8 | 4.7 | 5.0 |
| Mono/Macro (Patrolling) | 0.0 | 13.7 | 24.9 | 25.7 | 20.3 | 13.2 |
| Mono/Macro (*Ccr2*+) | 10.8 | 19.7 | 15.4 | 8.0 | 6.2 | 4.0 |
| Mono/Macro (*Mrc1*+) | 16.3 | 7.8 | 8.1 | 9.3 | 9.5 | 10.2 |
| Mono/Macro (*Cx3cr1*+) | 14.5 | 12.6 | 21.7 | 26.9 | 28.7 | 24.1 |
| Neutrophils | 9.3 | 32.0 | 9.7 | 2.4 | 2.4 | 2.9 |
| NK cells | 2.3 | 0.6 | 0.8 | 1.7 | 4.5 | 4.3 |
| T cells (3 CTs) | 12.5 | 1.9 | 2.1 | 3.9 | 10.3 | 20.0 |

**Supplementary Table 1: Determining the cell type composition in each day post injury.**
| Cell Types | % of Immune Cells |  |  |  |  |  |
| --- | --- | --- | --- | --- | --- | --- |
|  | 0 | 1 | 2 | 3.5 | 5 | 7 dpi |
| B cells | 14.6 | 0.6 | 0.4 | 0.3 | 0.5 | 4.6 |
| DC (*Cd209a*+) | 10.6 | 4.3 | 5.2 | 4.5 | 5.8 | 5.1 |
| DC (*Cd72*+) | 1.1 | 0.4 | 0.2 | 0.4 | 0.4 | 0.6 |
| DC (*Fscn1*+) | 0.0 | 0.3 | 0.3 | 0.6 | 1.5 | 1.5 |
| DC (*Xcr1*+) | 2.1 | 0.7 | 1.1 | 1.1 | 2.2 | 2.1 |
| Endo/Myeloid | 0.5 | 0.7 | 0.1 | 0.0 | 0.0 | 0.0 |
| Mono/Macro (Cycling) | 5.3 | 0.8 | 4.1 | 8.4 | 2.9 | 2.6 |
| Mono/Macro (*Cxcl10*+) | 0.3 | 4.0 | 5.9 | 6.8 | 4.7 | 5.0 |
| Mono/Macro (Patrolling) | 0.0 | 13.7 | 24.9 | 25.7 | 20.3 | 13.2 |
| Mono/Macro (*Ccr2*+) | 10.8 | 19.7 | 15.4 | 8.0 | 6.2 | 4.0 |
| Mono/Macro (*Mrc1*+) | 16.3 | 7.8 | 8.1 | 9.3 | 9.5 | 10.2 |
| Mono/Macro (*Cx3cr1*+) | 14.5 | 12.6 | 21.7 | 26.9 | 28.7 | 24.1 |
| Neutrophils | 9.3 | 32.0 | 9.7 | 2.4 | 2.4 | 2.9 |
| NK cells | 2.3 | 0.6 | 0.8 | 1.7 | 4.5 | 4.3 |
| T cells (*Cd4*+) | 0.3 | 0.2 | 0.4 | 0.5 | 0.7 | 1.0 |
| T cells (Cycling) | 0.6 | 0.0 | 0.1 | 0.3 | 0.6 | 1.1 |
| T cells (Non-cycling) | 11.6 | 1.7 | 1.6 | 3.1 | 9.0 | 17.8 |

**Supplementary Table 2:** **Cell type dynamics statistics summary.** For all cell types, listed the equation type used in the non-linear modeling, the unadjusted p-value, and the FDR adjusted p-value. The adjusted p-values that are significant (<0.05) are in red.

| Cell Type | Equation used in non-linear modeling | p-value | FDR-corrected p-value |
| --- | --- | --- | --- |
| B cells | Quadratic | 1.45E-03 | 4.06E-02 |
| Dendritic cells (*Cd209a+*) | Quadratic | 4.65E-02 | 1.00E+00 |
| Dendritic cells (*Cd72+*) | Quadratic | 3.60E-01 | 1.00E+00 |
| Dendritic cells (*Xcr1+*) | Quadratic | 2.02E-01 | 1.00E+00 |
| Endothelial and Myeloid cells | Quadratic | 7.66E-01 | 1.00E+00 |
| Endothelial cells (Capillary) | Quadratic | 7.27E-01 | 1.00E+00 |
| FAPs (Adipogenic) | Quadratic | 4.25E-01 | 1.00E+00 |
| FAPs (Stem) | Quadratic | 2.91E-02 | 8.14E-01 |
| Mono/Macro (*Cxcl10+*) | Quadratic | 8.30E-01 | 1.00E+00 |
| Myonuclei | Quadratic | 1.76E-01 | 1.00E+00 |
| Pericytes/SM | Quadratic | 2.62E-03 | 7.32E-02 |
| Schwann and Neural/Glial cells | Quadratic | 1.06E-01 | 1.00E+00 |
| T cells (*Cd4+*) | Quadratic | 3.36E-01 | 1.00E+00 |
| T cells (Cycling) | Quadratic | 4.37E-01 | 1.00E+00 |
| T cells (Non-cycling) | Quadratic | 7.06E-04 | 1.98E-02 |
| Tenocytes | Quadratic | 3.86E-02 | 1.00E+00 |
| Dendritic cells (*Fscn1+*) | Cubic | 5.65E-06 | 1.58E-04 |
| Endothelial cells (Artery) | Cubic | 1.08E-02 | 3.03E-01 |
| Endothelial cells (Vein) | Cubic | 7.71E-01 | 1.00E+00 |
| FAPs (Pro-remodeling) | Cubic | 5.06E-01 | 1.00E+00 |
| Mono/Macro (*Mrc1+*) | Cubic | 1.26E-08 | 3.52E-07 |
| Mono/Macro (*Cx3cr1+*) | Cubic | 1.92E-04 | 5.37E-03 |
| MuSCs and progenitors | Cubic | 4.10E-02 | 1.00E+00 |
| NK cells | Cubic | 8.17E-08 | 2.29E-06 |
| Mono/Macro (Cycling) | Quartic | 2.66E-05 | 7.46E-04 |
| Mono/Macro (Patrolling) | Quartic | 6.71E-06 | 1.88E-04 |
| Mono/Macro (*Ccr2+*) | Quartic | 1.00E-02 | 2.81E-01 |
| Neutrophils | Quartic | 9.47E-02 | 1.00E+00 |

## EXTENDED DATA FILES

**Extended Data File 1: Summary of single-cell RNA-sequencing samples in this study.**

**Extended Data File 2: Lists of senescence gene signatures used in this study.**

